# DeepD3, an Open Framework for Automated Quantification of Dendritic Spines

**DOI:** 10.1101/2023.02.01.526476

**Authors:** Martin H P Fernholz, Drago A Guggiana Nilo, Tobias Bonhoeffer, Andreas M Kist

**Author notes:** Corresponding author: Andreas M. Kist.

## Abstract

Dendritic spines are the seat of most excitatory synapses in the brain, and a cellular structure considered central to learning, memory, and activity-dependent plasticity. The quantification of dendritic spines from light microscopy data is usually performed by humans in a painstaking and error-prone process. We found that human-to-human variability is substantial (inter-rater reliability 82.2±6.4%), raising concerns about the reproducibility of experiments and the validity of using human-annotated ‘ground truth’ as an evaluation method for computational approaches of spine identification. To address this, we present DeepD3, an open deep learning-based framework to robustly quantify dendritic spines in microscopy data in a fully automated fashion. DeepD3’s neural networks have been trained on data from different sources and experimental conditions, annotated and segmented by multiple experts and they offer precise quantification of dendrites and dendritic spines. Importantly, these networks were validated in a number of datasets on varying acquisition modalities, species, anatomical locations and fluorescent indicators. The entire DeepD3 open framework, including the fully segmented training data, a benchmark that multiple experts have annotated, and the DeepD3 model zoo is fully available, addressing the lack of openly available datasets of dendritic spines while offering a ready-to-use, flexible, transparent, and reproducible spine quantification method.

## MAIN

Dendritic spines are small protrusions on dendrites. They constitute the postsynaptic part of most excitatory synapses in the brain. As such, dendritic spines have been postulated to act as the brain’s fundamental units of neuronal integration (Yuste and Denk, 1995) and the seat of information storage (e.g. Hofer and Bonhoeffer, 2010). Morphologically, dendritic spines are characterized by a bulbous head, which is connected to the dendrite via a thin spine neck (Yuste and Bonhoeffer, 2001).

Ongoing advances in microscopy have enabled researchers to obtain live images of dendritic spines, capturing fundamental mechanisms of synaptic plasticity, such as changes in spine size or quantity. However, the identification and quantification of dendritic spines are typically still done manually. This subjective, time-intensive task is further complicated by the limited spatial resolution common light microscopy techniques offer (Pfeiffer et al., 2018; Attardo et al., 2015). As a consequence, deriving a meaningful consensus across multiple raters is challenging. While the precise amount of inter-human variability in a spine identification task so far remains unclear, in comparable tasks, such as the identification of synapses or the classification of dendritic spines into morphological subtypes, inter-rater variability can reach levels of up to 30% (Graves et al., 2021; Rodriguez et al., 2008). This raises concerns about the reproducibility of the analysis for experiments involving the quantification of dendritic spines.

To address these concerns, various computational approaches of dendritic spine quantification have been described in the past (Extended Data Table 1) utilizing methods ranging from basic image thresholding to modern deep learning approaches (Koh et al., 2002; Dickstein et al., 2016; Xiao et al., 2018; Vidaurre-Gallart et al., 2022). These efforts improve the throughput of spine quantification and address the issue of reproducibility. However, they are typically tailored to perform well on data of a specific contrast and spatial resolution and hence perform inconsistently across datasets of different image qualities and modalities, hampering widespread use by the community (Xiao et al., 2018). Moreover, the validation of these approaches typically disregards human-to-human differences observed in manual spine identification, and instead compares the computational approach to a single human annotation (Koh et al.,2002; Xiao et al., 2018). Taken together, there are two main uncertainties: (1) human-to-human spine annotation variability and (2) data-related heterogeneity between experimental settings.

### Manual quantification of dendritic spines shows large inter- and intra-human variance

We reasoned that first a thorough investigation of human variability in spine quantification tasks is required to further improve automated approaches. To this end, a benchmark dataset was generated by two-photon imaging with a volume of 135.7 by 34.4 by 35.5 μm from the rat hippocampal CA1 region (94 × 94 × 500 nm voxel size). Next, multiple (n=7) experts manually annotated the center of mass of all dendritic spine heads in that dataset (Extended Data Fig. 1). The benchmark dataset, along with all annotations, is publicly available (see Data Availability) and constitutes, to the best of our knowledge, the first publicly available dataset of this kind.

We then determined the level of variability between human annotators by matching the manual annotations of dendritic spines using an unsupervised spatial clustering approach (see Online Methods). As expected from a subjective task such as spine annotations, rater-to-rater variability was considerable, resulting in an inter-rater-reliability (IRR) of 82.2±6.4% (Extended Data Fig. 1b). This is well in line with a previous report on a comparably subjective task (Graves et al., 2021, synapse identification; IRR: 72.3%) and underscored by the fact that less than 42.6% of all spines were found by all seven expert annotators in the benchmark dataset. Surprisingly, when annotators were tasked to identify dendritic spines in the same dataset several weeks later, variability was equally high for the same individual rating weeks apart (intra-rater-reliability: 87.5%, Extended Data Figure 1b).

Next, we asked multiple (n=3) experts to annotate the same benchmark dataset in a pixel-by-pixel manner into dendritic spines, dendrites, and background (see Online Methods). We found that while the individual experts agree on a qualitative level (Extended Data Figure 1a, d), quantitatively differences are sizeable: the Intersection over Union (IoU) score, a common measure of agreement in semantic segmentation tasks that ranges from 0 (no agreement) to 1 (perfect agreement), is on average 0.470±0.071 for dendrites and 0.423±0.094 for dendritic spines (Extended Data Fig. 1c). These results highlight two problems: First, our results call attention to the lack of reproducibility of studies involving spine quantification, since manual annotations are the current gold standard of spine counting and localization. Furthermore, our findings emphasize the necessity to involve multiple annotators when evaluating a dataset. The manual work of annotators already represents the main bottleneck in spine analysis pipelines, and the apparent need to perform this task multiple times further decreases the already low throughput of this process. Second, most automated methods of spine quantification have been using manual spine annotations as ground truth for both training and validation of the method. Here, too, the observed amount of inter- and intra-rater reliability suggests that multiple annotators are required for training and validation of automated means of spine quantification in order to minimize the subjective bias a single user introduces.

To address these observations, we have developed and report DeepD3, an open **Deep** Learning Framework for the **D**etection of **D**endritic Spines and **D**endrites. The DeepD3 framework employs training, validation, and benchmarking datasets that have been annotated by multiple experts, effectively addressing the variability observed among human annotators. The trained neural networks automatically perform semantic segmentation of dendrites and dendritic spines across microscopy data from different sources, offering a prompt solution to the slow, variable and error-prone manual spine annotation process. Furthermore, the emphasis on data heterogeneity during training and validation ensures that DeepD3 performs reliably across a range of data types, offering an open-source method for many applications, regardless of the provenance of the data.

### Semantic segmentation of dendrites and dendritic spines

To achieve this, we utilized supervised learning to adjust the network parameters of a deep convolutional neural network (see Online Methods), thereby optimizing its performance on this task (Figure 1a). To provide training data for supervised learning, dendritic spines and dendrites were human-expert-annotated in microscopy images with single-pixel precision (Figure 1b, see Online Methods). To account for user-dependent variability and ensure better generalizability of DeepD3, these data were generated by multiple (n=3) experts. This produced an extensive DeepD3 training dataset (Table 1), which consists of 3D microscopy image stacks of various resolutions, experimental data sources, imaging wavelengths, and microscopy modalities, all of which we provide as open data together with this study (see Data Availability). During the training of the deep neural network, we streamed single tiles of this training dataset, consisting of paired data containing the raw microscopy data, the binary dendritic spine segmentation mask, and the binary dendritic segmentation mask (Figure 1b, see Online Methods). Additionally, we applied data augmentations, such as rotation, flipping, blur, and Gaussian noise to these image tiles to generate more robust, generalized deep neural networks.

**Figure 1.**
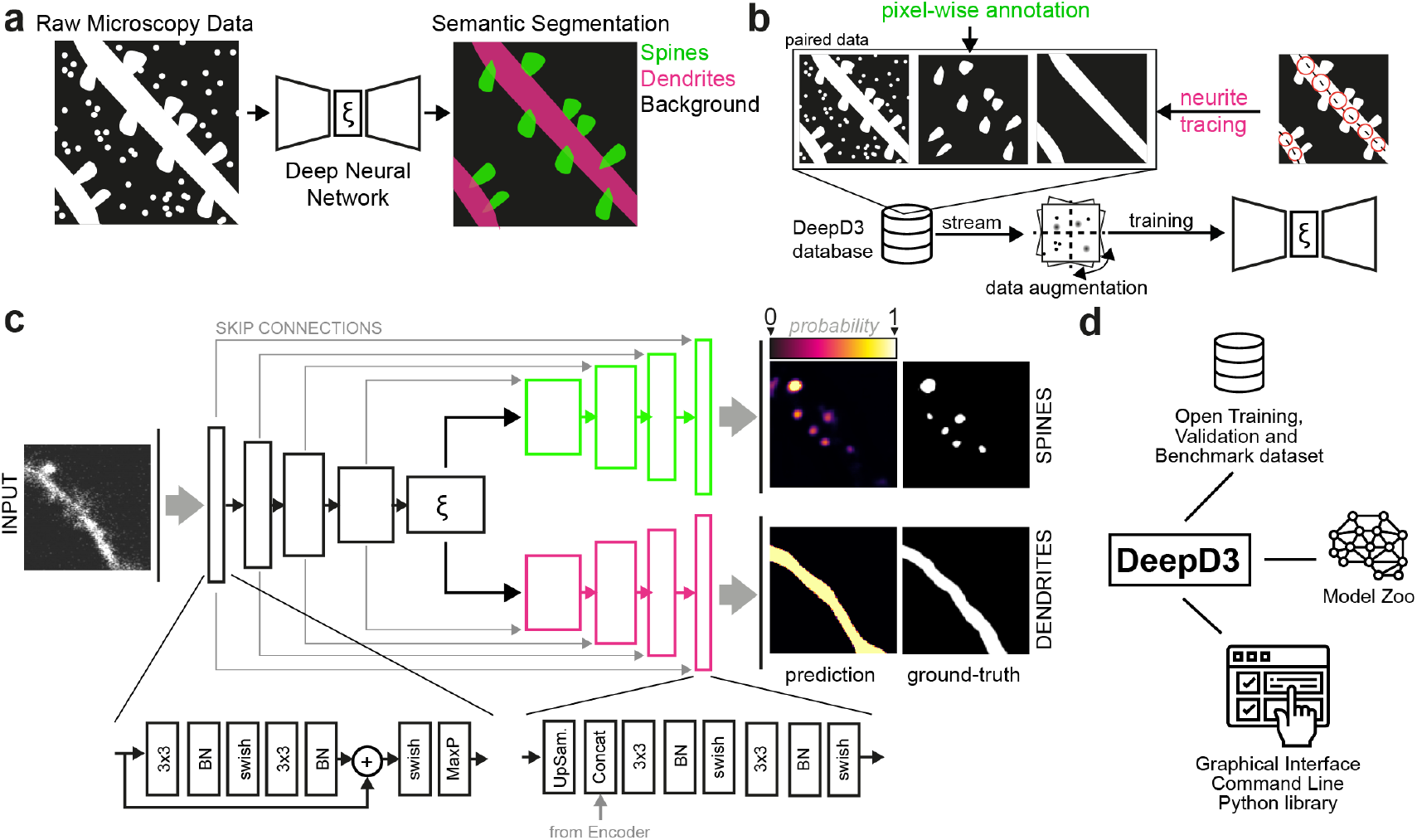
DeepD3 framework overview. **a**, Raw microscopy data (pictogram left) is used as input for a deep neural network (center) to semantically segment dendrites (magenta) and dendritic spines (green) against background (black; right). This color code will be used throughout the manuscript. **b**, DeepD3 database generation for paired ground-truth data. Before training, dendritic spines (top center) and dendrites (top right) are annotated in raw microscopy data (top left) using pixel-wise and semi-automatic tracing approaches (magenta circles in top far right image), respectively. During training, tiles from the DeepD3 database are streamed, dynamically augmented to increase variability, and fed into the DeepD3 training pipeline. **c**, The DeepD3 architecture features a dual-decoder structure that emerges from a common latent space *ξ* and receives skip connections from the encoder. Modules in the encoder are based on residual layers together with max pooling operations, whereas modules in the decoder contain upsampling operations, incorporate encoder input and use conventional convolutional layers. Example network input (left) and output (right) are shown as a microscopy image tile and a localization probability map ranging from 0 (background) to 1 (foreground). **d**, Overview of the DeepD3 open framework. DeepD3 consists of open datasets, a model zoo with training environment for custom neural networks, and a graphical user interface.

**Table 1.**
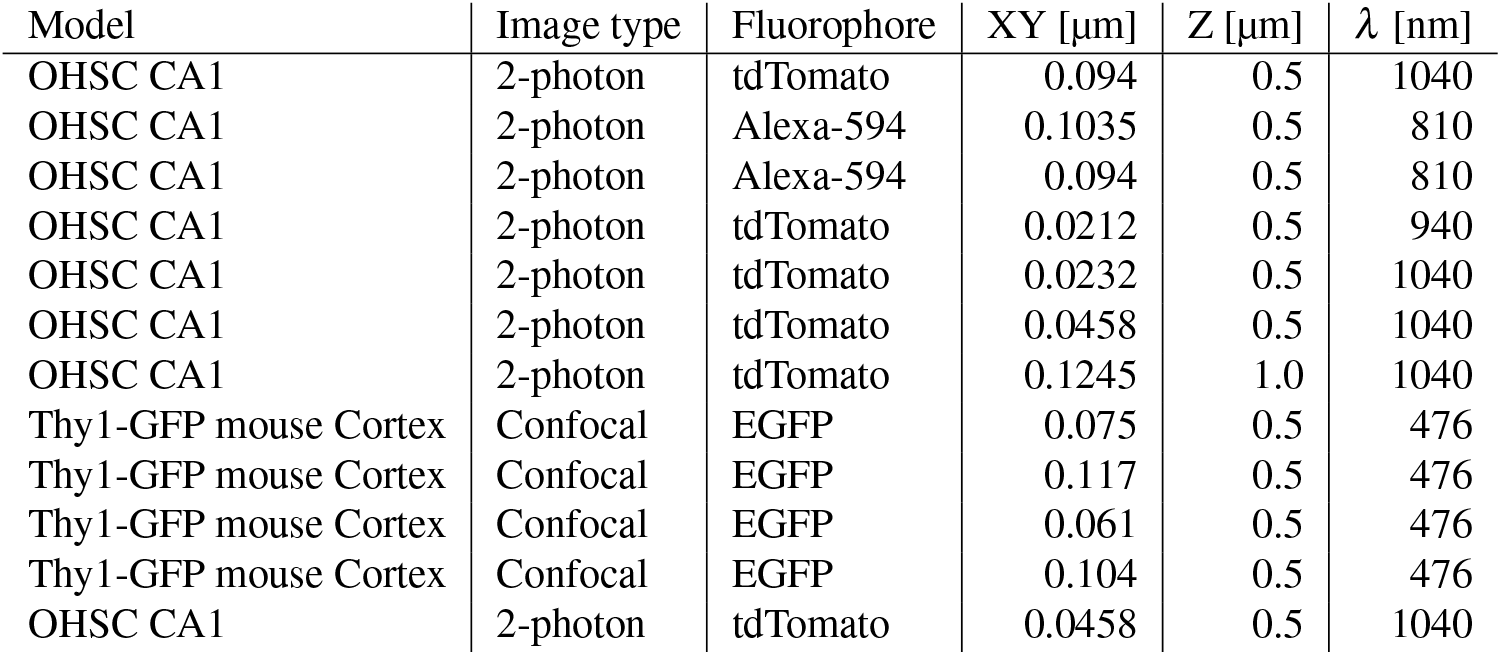
Overview of generated training data. Overview of generated training data, with model indicating the model organism and brain region (n.b.: all cells were pyramidal neurons), microscopy type, pixel size (resolution) in xy, and in z, imaging wavelength λ. All training data was generated in-house.

| Model | Image type | Fluorophore | XY [$\mu\text{m}$] | Z [$\mu\text{m}$] | $\lambda$ [nm] |
| --- | --- | --- | --- | --- | --- |
| OHSC CA1 | 2-photon | tdTomato | 0.094 | 0.5 | 1040 |
| OHSC CA1 | 2-photon | Alexa-594 | 0.1035 | 0.5 | 810 |
| OHSC CA1 | 2-photon | Alexa-594 | 0.094 | 0.5 | 810 |
| OHSC CA1 | 2-photon | tdTomato | 0.0212 | 0.5 | 940 |
| OHSC CA1 | 2-photon | tdTomato | 0.0232 | 0.5 | 1040 |
| OHSC CA1 | 2-photon | tdTomato | 0.0458 | 0.5 | 1040 |
| OHSC CA1 | 2-photon | tdTomato | 0.1245 | 1.0 | 1040 |
| Thy1-GFP mouse Cortex | Confocal | EGFP | 0.075 | 0.5 | 476 |
| Thy1-GFP mouse Cortex | Confocal | EGFP | 0.117 | 0.5 | 476 |
| Thy1-GFP mouse Cortex | Confocal | EGFP | 0.061 | 0.5 | 476 |
| Thy1-GFP mouse Cortex | Confocal | EGFP | 0.104 | 0.5 | 476 |
| OHSC CA1 | 2-photon | tdTomato | 0.0458 | 0.5 | 1040 |

To predict the presence of dendrites and dendritic spines in raw microscopy data, we observed that a custom two-decoder network (Figure 1c) inspired by the U-Net architecture (Ronneberger et al., 2015) performed better than an optimized vanilla U-Net (Extended Data Fig. 3). In the DeepD3 architecture, each decoder originates from a common latent space *ξ* that contains high-level image information extracted by the encoder. This allows independent optimization of dendrite and spine prediction without them interfering with each other. By adjusting the parameters of this dual-decoder architecture using the training dataset, DeepD3 networks successfully learned to segment dendrites and dendritic spines in microscopy image data (Extended Figure 3-5 and Figure 1c). By scaling the neural network, we show that DeepD3 adapts to changes in capacity, allowing fast and accurate variants (Extended Data Figures 4 and 5). For more details on the entire DeepD3 framework (Figure 1d), its elements (graphical user interface, user manual, datasets, model zoo, custom training environment and DeepD3 website), and its workflow, see the Supplementary Note, Table 1, Extended Data Figures 7-10, and Extended Data Tables 2-3.

### Validation on benchmark dataset

Given the observed amount of human-to-human variability (Extended Data Fig. 1), we concluded that the standard evaluation paradigm of automated spine detection methods is not suitable. I.e. comparing automated spine detection to a single manual ‘ground truth’ annotation and grouping spines into true positives, false positives, correct rejections, and false negatives seems inappropriate given that the ‘ground truth’ of a single annotator is so variable. Hence, we tasked DeepD3 to identify dendritic spines in the above-mentioned (DeepD3) benchmark dataset and compared the results to the performance of the seven human raters instead of a single one. Using the above-mentioned clustering approach, we were able to match spines between DeepD3 and the human raters and found that DeepD3 is able to recall most of the annotated dendritic spines by a given rater (91.6±2.8 %, Mann-Whitney U-test p¡0.01, see Figure 2b, right column). Inversely, human raters were less likely to recall spines that were identified by DeepD3 (76.0±4.5, see Figure 2b, bottom row). To verify human-like behavior of DeepD3, we utilized the matched spine annotations to investigate how many spines DeepD3 identifies relative to the spine cluster size, i.e. how many human annotators identified the same spine (Figure 2d). Spines that were identified by at least four of the seven human annotators were consistently also identified by DeepD3 (354 out of 369, 95.9%), while those that showed poorer agreement between human experts were also less likely to be found by DeepD3 (79 out of 122, i.e. 64.8%).

**Figure 2.**
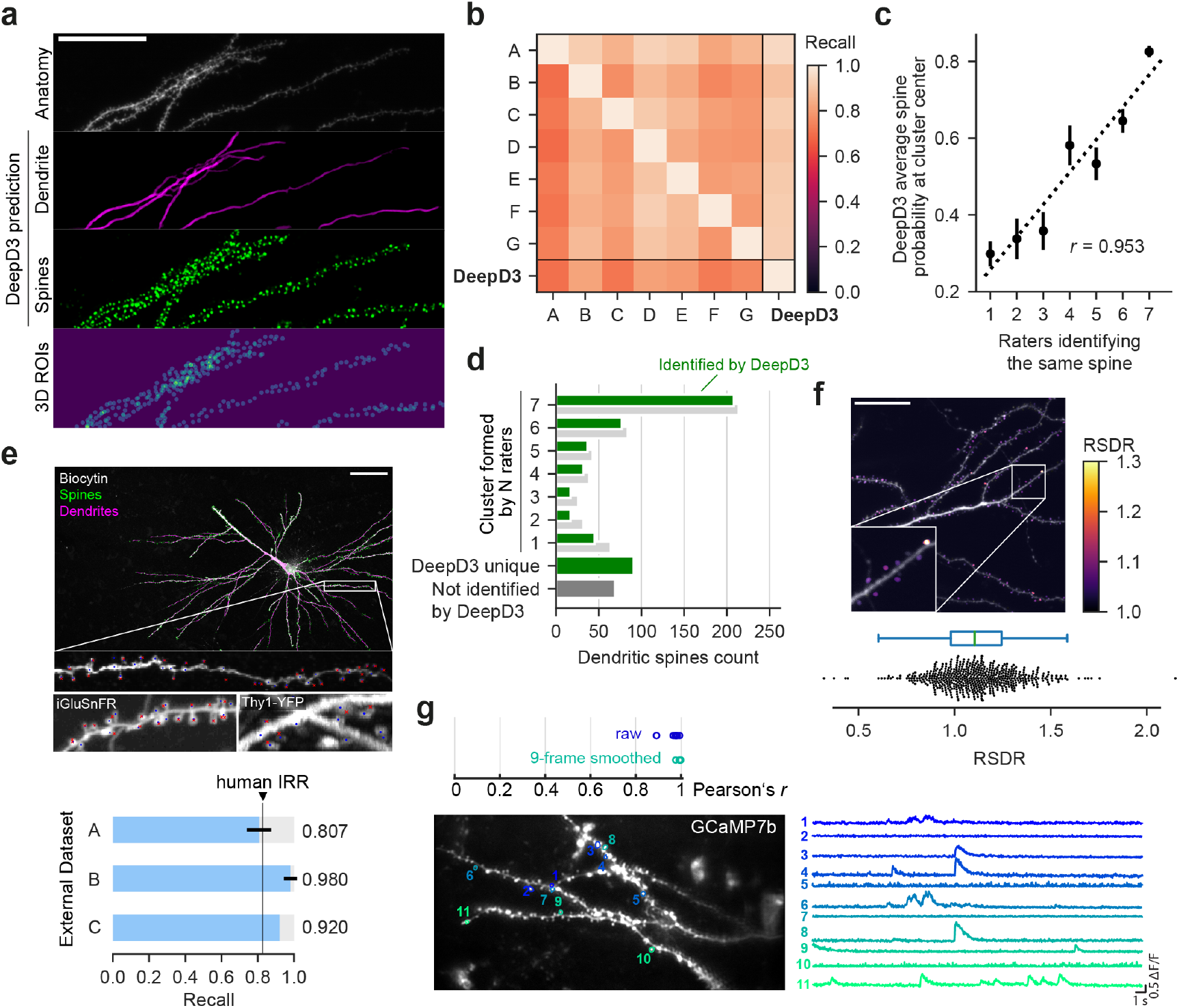
DeepD3 a versatile tool for quantification of dendritic spines in microscopy data. **a**, Maximum intensity projection of the benchmark dataset, a 3D image stack of dendrites and dendritic spines of CA1 pyramidal neuron of an organotypic hippocampal slice culture (raw data, top). DeepD3-generated prediction maps of dendrite (magenta) and dendritic spines (green). Segmented 3D ROIs using the spine prediction map (bottom). Scale bar indicates 50 μm. **b**, Inter-rater reliability of n=7 raters, who manually annotated the location of all dendritic spines in the benchmark dataset (see panel A). The matrix was generated by comparing rater pairs (y-axis = Rater 1, x-axis = Rater 2) using matched spine annotations (see Online Methods). The far right column indicates how many dendritic spines annotated by a given rater (y-axis, Rater 1) were identified by DeepD3 (x-axis, Rater 2). The bottom row indicates how many spines that were segmented by DeepD3 (here Rater 1) were also identified by a given human rater (here Rater 2). **c**, Linear correlation of the number of raters that identified a given spine and the average DeepD3 dendritic spine prediction probability at the center of the spine. Single points indicate the mean ±SEM, dashed line indicates the regression line. **d**, Frequency plot of the number of dendritic spines against the number of raters that identified a given spine (N={1,…,7}). Shown are the performances of DeepD3 (small green bars) and the raters (small gray bars). The bottom two bars plot the number of spines that were found by DeepD3 but none of the raters (single wide green bar) and those localized by a rater but not DeepD3 (single wide gray bar). **e**, Validation of DeepD3’s performance on three independently sourced and annotated datasets: In vivo iGluSnFR data was acquired in behaving mice using two-photon microscopy (dataset A; Kazemipour et al., 2019). In vivo Thy1-YFP data was acquired in behaving mice using two-photon microscopy (dataset B; Frank et al., 2018). In vitro image stacks of counter-stained Biocytin-filled neurons of human brain tissue were acquired using confocal microscopy (dataset C; Peng et al., 2015). Top: maximum intensity projections of example images of all three datasets with ground-truth annotation (red crosses) and DeepD3 3D ROI centroids (blue circles). Scale bar is 50 μm. Bottom: DeepD3 performance (recall) on the same datasets relative to the previously determined human IRR of the DeepD3 benchmark dataset (panels A and B). **f**, Utilization of DeepD3 for determining preferential localization of a nanobody against PSD-95 (Rimbault et al., 2021), tagged with mTurquoise2, to dendritic spines measured as the ratiometric spine-to-dendrite ratio (RSDR; see Online Methods). An RSDR value of >1 indicates preferential localization of the construct to the spine over the dendrite. Top: a maximum intensity projection of the raw data is overlayed with the RSDR of each DeepD3-generated spine ROI (purple to yellow, see color bar on the right). Scale bar is 50 μm. Bottom: box- and beeswarm plots of the RSDR measurements of all analyzed (N = 553) dendritic spines. **g**, Utilization of DeepD3’s dendritic spine segmentation for extracting calcium fluctuations of single dendritic spines using GCaMP7b. Bottom left: average projection of the analyzed calcium-imaging movie with DeepD3-generated spine ROI outlines in color and assigned numbers. Bottom right: calcium transients (ΔF/F_0_) of the outlined spines. Top: Pearsons’s correlation coefficient *r* of calcium transients of DeepD3-generated and manually generated spine ROIs using either raw data (blue) or 9-point moving averaged data (turquoise).

While, DeepD3 identified the vast majority of spines that had been annotated by at least one of the seven raters, it also generated some ‘unique’ spine annotations (i.e. those that could not be identified by any of the other raters) than the average rater (Figure 2d). We next investigated the origin of this ‘liberal annotation style’ of DeepD3 and whether it can be fine-tuned to the user’s needs. To that end, we utilized the fact that DeepD3 assigns prediction values to all pixels of an image as part of its workflow, with values ranging from 0 to 1, depending on the likelihood that DeepD3 deems a pixel to be part of a dendritic spine (see Online Methods for more details on the DeepD3 workflow). This spine prediction map was then utilized to compare the probability scores DeepD3 assigned to all dendritic spines to the number of human raters that identified a spine: not only is the probability DeepD3 assigns to a spine strongly correlated to how well humans detect that dendritic spine (Figure 2c, correlation coefficient *r* = 0.953), DeepD3’s prediction map also seems to reflect the distinct difference between those dendritic spines that have been identified by the majority of human experts versus those that have not (Figure 2c). Many of the user-defined hyper-parameters during DeepD3’s ROI generation process are implemented based on the spine prediction map (see Online Methods and Supplementary Note). Hence, and critically, fine-tuning such hyperparameters allows users to utilize DeepD3 as a more liberal or conservative tool with regard to spine ROI identification, depending on the user’s needs. We believe that an assessment of whether DeepD3 performs at, below or above the level of a human in a spine identification task is misguided, as the real ground truth for such an analysis is lacking. Instead, we show that DeepD3 by and large follows the spine identification behavior of human experts (Figure 2c,d) with the critical differences that hyper-parameters can be tuned to the requirements at hand in a reproducible and transparent manner while being considerably faster (Extended Data Figure 5a,b).

### Comparison to other automated methods

Using the segmentations of the benchmarking dataset provided by n=3 human experts, we next compared DeepD3 to state-of-the-art (semi-)automated methods for spine detection (Vidaurre-Gallart et al., 2022; Singh et al., 2017). DeepD3 (IoU for intersection: 0.474) outperformed a recently described fully automated deep learning method (Vidaurre-Gallart et al., 2022,; IoU for intersection: 0.278, Extended Data Figure 12), the semi-automated spine segmentation of the IMARIS platform (IoU of intersection: 0.343, Extended Data Figure 13), and a fully automated computational method (Singh et al., 2017, IoU for intersection: 0.422, Extended Data Figure 14). This indicates that DeepD3 is comparable or better than other state-of-the-art spine detection methods.

### Validation on diverse experimental settings

We next asked whether DeepD3 is also readily employable in other datasets. One key difference across experimental settings is the physical resolution of single pixels, measured as nanometers per pixel (nm/px). Past approaches of automating dendritic spine segmentation perform more poorly than reported in data with different or sometimes even similar physical resolution than their training dataset (Extended Data Figure 11), hampering widespread use in the community (see Extended Data Figures 12-14 and Xiao et al., 2018). To explore this, we trained the DeepD3 architecture in two independent manners: first, using a fixed resolution, i.e. all images of the training dataset were resized to comply with the same resolution of 94 nm/px in xy, and second, on a mix of all available resolutions in the training dataset (ranging from 23.2 to 124.5 nm/px in xy; see Table 1). Next, both of these trained neural nets (fixed and mixed) were tested against novel and rescaled (25-250 nm/px in xy) variants of the benchmark dataset. While both versions performed well in data with a spatial resolution close to 94 nm/px, the performance of the fixed neural net dropped when faced with images of vastly different spatial resolutions (Extended Data Figure 6a,c). The flexible neural net, on the other hand, performed spine segmentation much more reliably in a larger range of spatial resolutions (Extended Data Figure 6). Critically, this change together with other data augmentation strategies (see Online Methods) in the training procedure should enable the use of DeepD3 for data with a large range of image properties and thus considerably improve its applicability in the community.

To test this and extensively validate DeepD3, we gathered data from typical experimental paradigms in systems neuroscience (see Extended Data Table 2). Importantly, this data collection comprises high- and low-contrast anatomical data generated by different microscopy techniques (confocal, *ex vivo* and *in vivo* two-photon microscopy), using a variety of fluorescent indicators and dyes (YFP, iGluSnFR, tdTomato, GCaMP7b, biocytin + HRP + DAB) in different species (mouse, rat, human) and brain regions (retrosplenial cortex, hippocampus, frontal cortex; Figure 2e-g) Peng et al., 2015; Kazemipour et al., 2019; Frank et al., 2018). We determined the DeepD3 recall for spine localization across this collection and found that DeepD3 consistently performs at or above the human IRR (Figure 2e), despite large differences in the resolution of the data (67 to 240 nm/pixel). In contrast to other existing spine quantification methods, DeepD3 is therefore a readily employable tool for automated spine detection which is effective throughout a large variety of data sources with different resolutions, fluorophores, areas of origin and the like. Similar to other toolboxes, such as DeepLabCut (Mathis et al., 2018), we include the option to utilize transfer learning to improve performance even further.

Finally, we tried to extend DeepD3’s use cases to fluorescence extraction-based analyses by leveraging the fact that DeepD3 generates image segmentation for dendritic spines and dendrites. In particular, we asked whether we could capture the preferential localization of a major postsynaptic protein (PSD-95) to the dendritic spine using DeepD3-generated ROIs. Indeed, a nanobody against PSD-95 tagged with the fluorophore mTurquoise2 was found to be more prevalent in spines than in the dendritic arbor of single CA1 neurons (ratiometric spine-to-dendrite ratio >1, Figure 2f Rimbault et al., 2021, see Online Methods). In a separate experiment, DeepD3 was also found to be applicable to extract spontaneous synaptic transmission in the shape of calcium events at dendritic spines in two-photon calcium imaging data (Figure 2g): calcium traces of DeepD3-generated ROIs were strongly correlated with those generated by manually drawn spine ROIs (mean Pearson’s r = 0.97, see Figure 2g top, Extended Data Figure 15).

Taken together, we provide an open-source framework which we called DeepD3. In the course of generating the DeepD3 framework, we quantified user-dependent variability of manual spine annotations (available in the DeepD3 benchmark dataset). This led us to introduce a novel approach to evaluating spine detection performance of automated spine quantification methods: in light of the subjectivity of manual spine annotations and consequently the lack of true ‘‘ground truth”, we propose that an automated method’s performance should be evaluated against multiple annotations. DeepD3 performs well in such spine quantification tasks and has been validated for various data types. We were able to show that DeepD3 works in a human-like fashion but with much-improved speed and the benefit that spine quantification is performed in a transparent and reproducible manner. The entire DeepD3 framework is readily implemented in common microscopy analysis pipelines, providing an immediate end-to-end solution to the community. Lastly, by openly providing all training and benchmark datasets, a neural network model zoo, a method to train new networks, a GUI and batch execution function, as well as hosting the DeepD3 website to crowdsource and distribute annotated data and additions to the model zoo, we hope to have laid the foundation for a spine quantification tool, which will be useful for the community.

## ONLINE METHODS

### Sample preparation

#### Animals

All experiments were carried out in compliance with institutional guidelines of the Max Planck Society and of the local government (Regierung von Oberbayern). Wistar rats were housed under a 12 hour light-dark cycle with water and food available *ad libitum*. Pups (P4-9) were weaned and sacrificed to prepare organotypic hippocampal slice cultures the same day. Female Thy1-GFP mice were housed under a 12 hour light-dark cycle with water and food available *ad libitum*. One mouse (P100) was sacrificed to obtain the brain and generate brain slices.

#### DNA

The PSD-95-binding nanobody construct pCAG_Xph15-mTurquoise2-CCR5TC was generated via Gibson assembly by exchanging mNeonGreen (pCAG_Xph15-mNeonGreen-CCR5TC, Addgene 135533; Rimbault et al., 2021) with mTurquoise2. The sequence of the final construct was verified via PCR sequencing. pENN.AAV.CAG.tdTomato.WPRE.SV40 was obtained via Addgene (105554; Wilson lab, unpublished). pGP-AAV-syn-jGCaMP7b-WPRE was obtained via Addgene (104489, Dana et al., 2019).

#### Solutions

Sterile cortex buffer, sterile phosphate buffered saline (PBS), Cesium-based internal solution and K-gluconate-based intracellular recording solution were prepared as described earlier (Weiler et al., 2018). ACSF was prepared as described in Weiler et al. 2018 with the exception that calcium and magnesium concentrations were 3.7 mM and 0.15 mM, respectively. ACSF was supplied with D-serine (10 μM) and Trolox (1 mM). Culture medium for organotypic hippocampal slice cultures was prepared as described in Stoppini et al. 1991a and supplemented with penicillin (0.7 mM) and streptomycin (0.343 mM).

#### Organotypic Hippocampal Slice Culture Preparation

Organotypic hippocampal slice cultures (OHSCs) were prepared from Wistar rats on P4-9 (day of birth = P0) by cutting 400 μm transverse section as previously described (Stoppini et al., 1991b). OHSCs were kept in an incubator at 35°C with 5 % CO_2_ enriched atmosphere. Medium (50 % volume of each well) was exchanged twice a week.

#### Single-Cell Electroporation

To express DNA constructs in single hippocampal CA1 neurons (DIV 10-18), single-cell electroporation (SCE) was performed, as described previously (Judkewitz et al., 2009). Utilized constructs were pENN-AAV-CAG-tdTomato-WPRE-SV40 and either pGP-AAV-syn-jGCaMP7b-WPRE or pCAG_Xph15-mTurquoise2-CCR5TC, diluted to a total concentration of 100 ng/ml in K-gluconate-based intracellular solution. Constructs were given 2-8 days to express while slice cultures remained in the incubator.

#### Immunohistochemistry

A Thy1-GFP mouse brain (P100) was sliced into 300 μm thick coronal brain slices, which were subsequently brain cleared using RapiClear 1.47 for ~3 h. Subsequently, slices were embedded in RapiClear solution and placed on a coverslip with a custom-made 300 μm spacer to prevent squeezing of the slice.

#### Electrophysiology

To obtain structural imaging data of dendrites and spines with different signal-to-noise ratios to the used genetically encoded structural markers (tdTomato/EGFP), patch-clamp recordings were performed as described earlier (Bauer et al., 2021). To allow for sufficient filling of pyramidal CA1 neurons, at least 15 minutes separated break-in time and the beginning of structural imaging.

#### Pharmacology

Imaging of rat OHSCs was performed in the presence of voltage-gated potassium and sodium channel inhibitors (4-AP and TTX, respectively; both obtained from Tocris, USA).

#### Two photon setup

A custom-made two-photon imaging setup was utilized for imaging dendrites and dendritic spines in OHSCs. An 80 MHz pulsed femtosecond Ti:sapphire laser (MaiTai eHP, Spectra-Physics, USA) was used as excitation source for two-photon (2p) imaging. An electro-optical modulator (Pockels cell; Conoptics, USA, Model 350-80) was utilized for turnaround blanking and control over 2p excitation intensity. A galvo-resonant scan system (8 kHz) was utilized for bidirectional raster-scanning. A total of four photo-multiplier tube detection modules (PMTs) captured emitting photons in two separate detection paths: two PMTs were placed epidirectionally (via the objective) and two transdirectionally to the objective (via a 1.4 NA oil immersion condenser; Thorlabs, Germany). Photons were spectrally separated (560nm dichroic beam splitters) and filtered (red/green imaging: 525-50nm and 607-70nm bandpass filters; blue-shifted fluorophores: 510-84nm bandpass filter and 6.0ND filter) according to use before hitting the PMTs, such that always one PMT of each detection path captured photons of the same wavelength. Images were obtained using a 60x, 1.1 NA water immersion objective (Olympus, Japan).

Calcium imaging was performed using volumetric two-photon imaging, as described earlier (Lu et al.,2017). The desired light pattern was achieved using a spatial light modulator (SLM, XY-series, Meadowlark, USA) and produced a z-elongated point spread function (FWHM ~ 14 μm), which allowed for effective stimulation of a volume.

### Data acquisition

Structural and Xph-15-mTurquoise-based images of rat CA1 pyramidal neurons in OHSCs were acquired at different spatial resolutions (see Table 1), with a 60x 1.1 NA water immersion objective (Olympus, Japan). Image acquisition was usually performed at a resolution of 2048 × 2048 px (7.3 Hz), a step size of 0.5 μm (using a piezoelectric z-scanner, Physik Instrumente, Germany), while frame averaging 50 frames online. Excitation wavelengths for TdTomato, Alexa-594 and mTurquoise2 were 1040 nm, 810 nm, and 860 nm, respectively. Average laser power under the objective was kept below 15 mW.

Calcium images were acquired at a temporal resolution of 30 Hz with a spatial resolution of 1024 × 512 px using extended depth-of-field Bessel beam imaging (see above). GCaMP7b was imaged using an excitation wavelength of 940 nm. Two-photon images were acquired using ScanImage r4.2 (Mathworks, USA).

Cleared organotypic hippocampal slice cultures (OHSCs) were imaged using a confocal microscope (Sp8, Leica, Germany) at voxel sizes of 0.061-0.117 × 0.061-0.117 × 0.5 μm equipped with an argon-ion laser (476 nm). Images were acquired through an HC PL APO L 20x/0.75 IMM CORR CS2 objective (Leica, Germany), scanning bidirectionally at 600 Hz. Emitting photons were captured via a PMT (488 nm - 738 nm). Images were captured with a resolution of 1024 × 1024 pixel, online averaging 20 frames per z level.

Structural images (iGluSnFR) of L1 and L2/3 neurons in mouse V1 were obtained with permission from the authors and generated as previously described (Kazemipour et al., 2019). Structural images (YFP) of L5 neurons of mouse retrosplenial cortex were obtained with permission from the authors and generated as previously described (Frank et al., 2018). Structural images of human cortical neuron were obtained via the BigNeuron project (gold166 dataset, http://bigneuron.org) of the Allen Brain Institute (Peng et al., 2015; Chen et al., 2017; Manubens-Gil et al., 2022).

### Data processing

All two-photon images of rat CA1 pyramidal neurons were deinterlaced using custom-written Matlab code. Deinterlaced structural and PSD-95-nanobody images of rat CA1 pyramidal neurons were first manually registered using custom-written Matlab code and subsequently registered using the Computational Morphometry Toolkit (CMTK, https://www.nitrc.org/projects/cmtk/). ROIs of dendritic spines and dendrites were generated using DeepD3. ROI outlines were utilized to extract average raw fluorescence values of dendrites and spines in 3D from the two registered image stacks (tdTomato, structural; mTurquoise2, PSD-95-nanobody). To quantify the localization preference of the nanobody Xph-15 (Rimbault et al.,2021), the ratiometric spine-to-dendrite-ratio (RSDR) was utilized, comparing the ratiometric expression levels of Xph-15-mTurquoise2 and tdTomato of each spine to that of the entire dendritic arbor in the image:

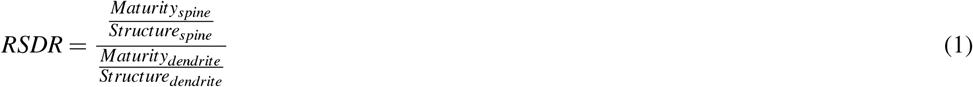

Where Maturity_spine_ and Structure_spine_ are the average raw fluorescence values of all voxels within a given spine ROI of the maturity and structural images, respectively. Maturity_dendrite_ and Structure_dendrite_ are the mean raw fluorescence values of all voxels that had been labeled as dendrite, extracted from the maturity and structural images, respectively. RSDR values of above one indicate preferential localization of the PSD-95-nanobody to a dendritic spine over the dendrite.

Deinterlaced calcium images were registered via cross-correlation using custom-written Matlab code. ROIs of dendritic spines were either generated using DeepD3 or manually performed using PiPrA (Gómez et al., 2020). ROI outlines were utilized to extract average raw fluorescence values of dendrites and spines per frame. Using these time series, normalized fluorescence fluctuations were computed:

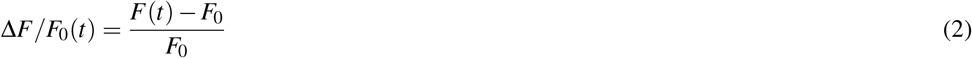

where *F* is the mean fluorescence of the green channel of a given ROI at timepoint *t*, and *F*_0_ is the mean raw fluorescence of that same channel during baseline (20 frames without spontaneous calcium events).

The benchmarking dataset was filtered using a Gaussian filter (sigma = 2) to improve the segmentation performance of IMARIS (Bitplane, USA).

### Manual annotation of dendritic spines

To identify the location of dendritic spines in three-dimensional microscopy stacks, human experts were tasked to manually annotate the (x,y,z) position of the dendritic spine that was closest to the center of mass of the spine head. Experts were tasked to annotate all dendritic spines and utilized the ImageJ ROI Manager for this annotation task.

### Manual segmentation of dendritic spines

To segment dendritic spines in images, a custom open-source graphical user interface (PiPrA; Gómez et al., 2020) was utilized to perform pixel-wise annotations for spine labeling. The resulting binary segmentation masks were used as input data for training of model networks (see below) or to extract average raw fluorescence values from images.

### Current spine segmentation methods

Segmentation of a smoothed version of the benchmarking dataset was performed with IMARIS 9.9.0 (Bitplane, USA) using the following parameters: dendrites were semi-automatically traced using the filament tracer in the AutoPath setting (threshold: 1.31) and spine seed points generated (0.25-2.5 μm) and filled (threshold: 0.82). The resulting surface was exported in .wrl format, subsequently converted to .stl format using MeshLab v2022.02 (Ranzuglia et al., 2013), and finally converted to binary .tif files using custom-written Matlab code. As a consequence, spine and dendrite signal could no longer be differentiated, which is why IoU scores were calculated on the union of both segmentation sources (see also Extended Data Figure 12).

The benchmarking dataset was also segmented using two recently published automated spine detection methods (Vidaurre-Gallart et al., 2022; Singh et al., 2017). NB: centerline extraction and hence differentiation of spine and dendrite signal did not work in our hands using the code provided by Singh et al., (2017) reducing comparability to IoU scores of both spine and dendrite signal.

### Training data generation

We manually labeled a collection of in-house acquired data to generate the DeepD3 training dataset (1). Manual pixel-precise annotations of dendritic spines were generated as described above. Dendrites were traced in three dimensions using NeuTube (Feng et al., 2015). Traced dendrites were saved in the SWC file format. We used a Python-based custom-written toolbox to generate a TIFF stack with the same dimension as the respective stack based on the SWC annotation files by drawing interpolated spheres between the anchor points traced and defined by NeuTube. The code is openly available together with the DeepD3 framework.

### Benchmark dataset generation

The benchmark dataset consists of a z-stack that was manually annotated by seven independent experts (see manual annotation of dendritic spines) to compute the inter-rater reliability. We used DBSCAN to cluster the point cloud (the pooled seven center-of-mass annotations) into individual clusters to identify annotations of the different human experts that pertain to individual dendritic spines. After DBSCAN’s initial cluster creation, we used *a priori* knowledge to adjust the clusters accordingly: (i) we removed points that were spatially too distant from the cluster center (0.85 μm in x and y, 2.5 μm in z), (ii) we split clusters that have more than seven annotations using K-Means (n_clusters = 2), (iii) we merged clusters that were closer than 0.85 μm in x and y and closer than 2.5 μm in z, but only if the new cluster did not exceed a total of seven annotations. This cluster map is used to assess inter- and intra-rater reliability and to evaluate the performance of DeepD3 in relation to human annotators (Figure 2b,d). Additionally, two of the seven experts annotated the full z-stack twice to also determine intra-rater reliability using the same clustering approach. The two annotation rounds were separated by at least 14 days to prevent carry-over effects from the first annotation. Further, three experts labeled the dataset in a pixel-precise fashion using PiPrA and NeuTube to assess inter-rater reliability on a segmentation level (Extended Data Figure 1c). All of this data is available on zenodo: https://doi.org/10.5281/zenodo.7590772.

### Deep neural networks

The DeepD3architecture is based on an encoder-decoder neural architecture that condenses information to the bottleneck latent space ξ, which provides a high-level abstract embedding of the input image. Our architecture contains a dual-decoder structure for spines and dendrites, respectively, and it is similar in its main concept to the U-Net architecture (Ronneberger et al., 2015) that has seen widespread attention in the biomedical domain (Falk et al., 2019; Isensee et al., 2021; Stringer et al., 2021). In detail, we use 3 × 3 kernel sizes, Batch Normalization layers (Ioffe and Szegedy, 2015) before activation, and the swish activation function (Ramachandran et al., 2017) for all convolutional layers (eq. 4), except the last one, where the logistic function (eq. 3) is used to ensure a dendrite/spine probability between 0 and 1. We further utilize residual connections He et al. (2016).

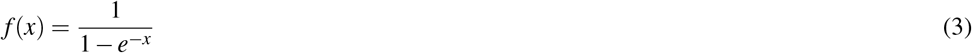

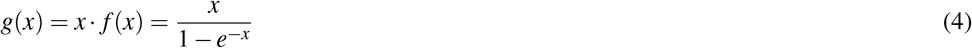

We optimize the DeepD3architecture using two losses, the Dice loss (eq. 5) and the mean squared error (eq. 6), for the semantic segmentation of dendrites and spines, respectively.

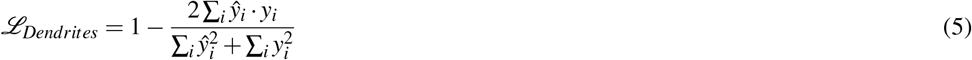

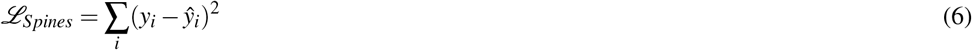

Images are provided with a single channel, where image intensities are by default linearly rescaled to -1 (minimum Intensity) and 1 (maximum Intensity). We trained **DeepD3** on two-dimensional 128 × 128 px tiles dynamically streamed from z-stacks (see 1b) and a learning rate starting at 0.5 · 10^−3^ for 15 epochs and then with an exponential decay with a time constant of 0.1. The network was optimized using the Adam optimizer (Kingma and Ba, 2014). Networks were set up in TensorFlow using the high-level Keras package (Abadi et al., 2016).

We used realistic just-in-time data augmentation (Figure 1b) to increase our training dataset. These included random rotation (90 °and *r* ∈ [−10,10]) and flipping, Gaussian noise, brightness and contrast changes. We provide the training dataset openly on Zenodo (see Data Availability).

### Prediction postprocessing

We first removed dendritic prediction segments in 2D and 3D that were too small for a valid patch using binary thresholded 3D connected component analysis. Next, we dilated this cleaned dendritic prediction map such that we included all spine predictions that were close to a dendrite, therefore implicitly incorporating a distance-to-dendrite metric ensuring that a spine is next to a dendrite. In general, these postprocessing options are optional and can be adjusted according to the special use case.

### ROI detection

Regions of interest (ROIs) are identified spines in either 2D or 3D using different methods. We either use a custom flood-filling paradigm or perform threshold-based 3D connected component analysis (Extended Data Figure 9). In general, each successfully identified ROI complies with a given set of properties. For custom flood filling: (1) spine prediction above user-set threshold to determine a seed pixel to create a novel ROI, (2) maximum euclidean distance to seed pixel, (3) relation to seed pixel. For 3D connected component analysis, spine prediction needs to be above a user-set threshold to determine the area of a given spine. Both methods set further requirements to keep an ROI: (1) minimum ROI size, (2) maximum ROI size, (3) minimum planes span, (4) implicit or explicit maximum distance to dendrite (see also section **Prediction postprocessing**). We measure the distances in μm using the user-defined resolution in **x**, *y*, and *z* of the stack. The settings can be combined if desired.

### Graphical user interfaces

As a companion to DeepD3, we provide two graphical user interfaces (GUIs): one, to prepare training data according to custom needs (Extended Data Fig. 10), and a second GUI, to infer dendrite and spine labels from a loaded dataset and to fully automatically create a set of two- or three-dimensional ROIs (Extended Data Fig. 7-9). Both GUIs are written in Python 3.7+ and are based on PyQt5 as graphical interface and pyqtgraph for plotting. We further allow the export of predictions to *.tif files and created ROIs to the ImageJ/FIJI-specific format to maximize interoperability with existing ecosystems. Further information is provided in the Supplementary Note.

### Evaluation

We used private and public datasets for evaluating DeepD3. We focused on multiple species, namely rat, mouse and human, as well as imaging modalities, namely *ex vivo* and *in vivo* two-photon and confocal microscopy, across several brain areas including V1 and the hippocampus. To quantify performance, we computed the recall metric (eq. 7). TPs are the true positive, and FNs the false negative detected dendritic spines. To evaluate the pixel-wise agreement across segmentation masks, we use the Intersection over Union (IoU) score (eq. 8). When comparing the performance of DeepD3 to other state-of-the-art spine identification methods, we use a combination of all three pixel-precise, human-annotated segmentation masks (provided by raters U, V and W): the intersection of all raters (*U* ∩ *V* ∩ *W*), the union of all raters (*U* ∪ *V* ∪ *W*) and the individual segmentation mask (Extended Data Figure 12).

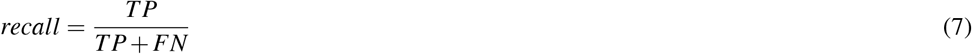

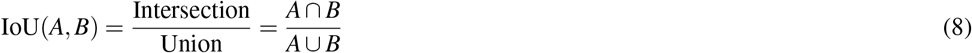

## CODE AVAILABILITY

All code is openly available at https://github.com/ankilab/DeepD3.

## DATA AVAILABILITY

The training data and pre-trained deep neural networks are available at https://doi.org/10.5281/zenodo.7590772 and https://doi.org/10.5281/zenodo.7590704, respectively. All data can be accessed through the project website at https://deepd3.forschung.fau.de/.

## ACKNOWLEDGMENTS

We thank Shan Huang, Alcino Silva, Kaspar Podgorski, Filip Tomaska, and Ruth Benavides-Piccione for sharing spine microscopy data and annotations. We thank Volker Staiger, Claudia Huber, Hiranmay Girish Joag, Danielle Paynter, Dominik Lindner, Joel Bauer, Adrianna Zgraj, and Sascha-Alexander Heye for manual annotation of dendritic spines. We thank the Matthieu Sainlos, James M. Wilson, Karl Deisseroth, and Douglas Kim for kindly providing DNA constructs. We thank Volker Staiger, Claudia Huber, Tobias Rose, and Robert Kasper for technical assistance. We thank Joergen Kornfeld and Joel Bauer with for helpful suggestions and critical comments on the manuscript. AMK was funded by the High-Tech Agenda Bavaria. MHPF was funded by the Boehringer Ingelheim Fonds PhD stipend. MHPF, DGN and TB were funded by the Max Planck society.

## REFERENCES

Abadi, M., Barham, P., Chen, J., Chen, Z., Davis, A., Dean, J., Devin, M., Ghemawat, S., Irving, G., Isard, M., et al. (2016). Tensorflow: A system for large-scale machine learning. In 12th {USENIX} symposium on operating systems design and implementation ({OSDI} 16), pages 265–283.

Attardo, A., Fitzgerald, J. E., and Schnitzer, M. J. (2015). Impermanence of dendritic spines in live adult ca1 hippocampus. Nature, 523(7562):592–596.

Bauer, J., Weiler, S., Fernholz, M. H., Laubender, D., Scheuss, V., Hübener, M., Bonhoeffer, T., and Rose, T. (2021). Limited functional convergence of eye-specific inputs in the retinogeniculate pathway of the mouse. Neuron, 109(15):2457–2468.

Chen, H., Iascone, D. M., da Costa, N. M., Lein, E. S., Liu, T., and Peng, H. (2017). Fast assembling of neuron fragments in serial 3d sections. Brain informatics, 4(3):183–186.

Dana, H., Sun, Y., Mohar, B., Hulse, B. K., Kerlin, A. M., Hasseman, J. P., Tsegaye, G., Tsang, A., Wong, A., Patel, R., et al. (2019). High-performance calcium sensors for imaging activity in neuronal populations and microcompartments. Nature methods, 16(7):649–657.

Dickstein, D. L., Dickstein, D. R., Janssen, W. G., Hof, P. R., Glaser, J. R., Rodriguez, A., O’Connor, N., Angstman, P., and Tappan, S. J. (2016). Automatic dendritic spine quantification from confocal data with neurolucida 360. Current protocols in neuroscience, 77(1):1–27.

Falk, T., Mai, D., Bensch, R., Çiçek, Ö., Abdulkadir, A., Marrakchi, Y., Böhm, A., Deubner, J., Jäckel, Z., Seiwald, K., et al. (2019). U-net: deep learning for cell counting, detection, and morphometry. Nature methods, 16(1):67–70.

Feng, L., Zhao, T., and Kim, J. (2015). neutube 1.0: a new design for efficient neuron reconstruction software based on the swc format. eneuro, 2(1).

Frank, A. C., Huang, S., Zhou, M., Gdalyahu, A., Kastellakis, G., Silva, T. K., Lu, E., Wen, X., Poirazi, P., Trachtenberg, J. T., et al. (2018). Hotspots of dendritic spine turnover facilitate clustered spine addition and learning and memory. Nature communications, 9(1):1–11.

Gómez, P., Kist, A. M., Schlegel, P., Berry, D. A., Chhetri, D. K., Dürr, S., Echternach, M., Johnson, A. M., Kniesburges, S., Kunduk, M., et al. (2020). Bagls, a multihospital benchmark for automatic glottis segmentation. Scientific data, 7(1):186.

Graves, A. R., Roth, R. H., Tan, H. L., Zhu, Q., Bygrave, A. M., Lopez-Ortega, E., Hong, I., Spiegel, A. C., Johnson, R. C., Vogelstein, J. T., et al. (2021). Visualizing synaptic plasticity in vivo by large-scale imaging of endogenous ampa receptors. Elife, 10:e66809.

He, K., Zhang, X., Ren, S., and Sun, J. (2016). Deep residual learning for image recognition. In Proceedings of the IEEE conference on computer vision and pattern recognition, pages 770–778.

Hofer, S. B. and Bonhoeffer, T. (2010). Dendritic spines: the stuff that memories are made of? Current biology, 20(4):R157–R159.

Ioffe, S. and Szegedy, C. (2015). Batch normalization: Accelerating deep network training by reducing internal covariate shift. In International conference on machine learning, pages 448–456. PMLR.

Isensee, F., Jaeger, P. F., Kohl, S. A., Petersen, J., and Maier-Hein, K. H. (2021). nnu-net: a self-configuring method for deep learning-based biomedical image segmentation. Nature methods, 18(2):203–211.

Judkewitz, B., Rizzi, M., Kitamura, K., and Häusser, M. (2009). Targeted single-cell electroporation of mammalian neurons in vivo. Nature protocols, 4(6):862–869.

Kazemipour, A., Novak, O., Flickinger, D., Marvin, J. S., Abdelfattah, A. S., King, J., Borden, P. M., Kim, J. J., Al-Abdullatif, S. H., Deal, P. E., et al. (2019). Kilohertz frame-rate two-photon tomography. Nature methods, 16(8):778–786.

Kingma, D. P. and Ba, J. (2014). Adam: A method for stochastic optimization. arXiv preprint arXiv:1412.6980.

Koh, I. Y., Lindquist, W. B., Zito, K., Nimchinsky, E. A., and Svoboda, K. (2002). An image analysis algorithm for dendritic spines. Neural computation, 14(6):1283–1310.

Lu, R., Sun, W., Liang, Y., Kerlin, A., Bierfeld, J., Seelig, J. D., Wilson, D. E., Scholl, B., Mohar, B., Tanimoto, M., et al. (2017). Video-rate volumetric functional imaging of the brain at synaptic resolution. Nature neuroscience, 20(4):620–628.

Manubens-Gil, L., Zhou, Z., Chen, H., Ramanathan, A., Liu, X., Liu, Y., Bria, A., Gillette, T., Ruan, Z., Yang, J., et al. (2022). Bigneuron: A resource to benchmark and predict best-performing algorithms for automated reconstruction of neuronal morphology. bioRxiv.

Mathis, A., Mamidanna, P., Cury, K. M., Abe, T., Murthy, V. N., Mathis, M. W., and Bethge, M. (2018). Deeplabcut: markerless pose estimation of user-defined body parts with deep learning. Nature neuroscience, 21(9):1281–1289.

Peng, H., Hawrylycz, M., Roskams, J., Hill, S., Spruston, N., Meijering, E., and Ascoli, G. A. (2015). Bigneuron: large-scale 3d neuron reconstruction from optical microscopy images. Neuron, 87(2):252–256.

Pfeiffer, T., Poll, S., Bancelin, S., Angibaud, J., Inavalli, V. K., Keppler, K., Mittag, M., Fuhrmann, M., and Nägerl, U. V. (2018). Chronic 2p-sted imaging reveals high turnover of dendritic spines in the hippocampus in vivo. Elife, 7:e34700.

Ramachandran, P., Zoph, B., and Le, Q. V. (2017). Searching for activation functions. arXiv preprint arXiv:1710.05941.

Ranzuglia, G., Callieri, M., Dellepiane, M., Cignoni, P., and Scopigno, R. (2013). Meshlab as a complete tool for the integration of photos and color with high resolution 3d geometry data. In CAA 2012 Conference Proceedings, pages 406–416. Pallas Publications - Amsterdam University Press (AUP).

Rimbault, C., Breillat, C., Compans, B., Toulmé, E., Vicente, F. N., Fernandez-Monreal, M., Mascalchi, P., Genuer, C., Puente-Muñoz, V., Gauthereau, I., et al. (2021). Engineering paralog-specific psd-95 synthetic binders as potent and minimally invasive imaging probes. BioRxiv.

Rodriguez, A., Ehlenberger, D. B., Dickstein, D. L., Hof, P. R., and Wearne, S. L. (2008). Automated three-dimensional detection and shape classification of dendritic spines from fluorescence microscopy images. PloS one, 3(4):e1997.

Rohlfing, T. and Maurer, C. R. (2003). Nonrigid image registration in shared-memory multiprocessor environments with application to brains, breasts, and bees. IEEE transactions on information technology in biomedicine, 7(1):16–25.

Ronneberger, O., Fischer, P., and Brox, T. (2015). U-net: Convolutional networks for biomedical image segmentation. In International Conference on Medical image computing and computer-assisted intervention, pages 234–241. Springer.

Singh, P. K., Hernandez-Herrera, P., Labate, D., and Papadakis, M. (2017). Automated 3-d detection of dendritic spines from in vivo two-photon image stacks. Neuroinformatics, 15(4):303–319.

Stoppini, L., Buchs, P.-A., and Muller, D. (1991a). A simple method for organotypic cultures of nervous tissue. Journal of neuroscience methods, 37(2):173–182.

Stoppini, L., Buchs, P.-A., and Muller, D. (1991b). A simple method for organotypic cultures of nervous tissue. Journal of Neuroscience Methods, 37(2):173–182.

Stringer, C., Wang, T., Michaelos, M., and Pachitariu, M. (2021). Cellpose: a generalist algorithm for cellular segmentation. Nature Methods, 18(1):100–106.

Vidaurre-Gallart, I., Fernaud-Espinosa, I., Cosmin-Toader, N., Talavera-Martínez, L., Martin-Abadal, M., Benavides-Piccione, R., Gonzalez-Cid, Y., Pastor, L., DeFelipe, J., and García-Lorenzo, M. (2022). A deep learning-based workflow for dendritic spine segmentation. Frontiers in neuroanatomy, 16:817903–817903.

Weiler, S., Bauer, J., Hübener, M., Bonhoeffer, T., Rose, T., and Scheuss, V. (2018). High-yield in vitro recordings from neurons functionally characterized in vivo. Nature protocols, 13(6):1275–1293.

Xiao, X., Djurisic, M., Hoogi, A., Sapp, R. W., Shatz, C. J., and Rubin, D. L. (2018). Automated dendritic spine detection using convolutional neural networks on maximum intensity projected microscopic volumes. Journal of neuroscience methods, 309:25–34.

Yuste, R. and Bonhoeffer, T. (2001). Morphological changes in dendritic spines associated with long-term synaptic plasticity. Annual review of neuroscience, 24(1):1071–1089.

Yuste, R. and Denk, W. (1995). Dendritic spines as basic functional units of neuronal integration. Nature, 375(6533):682–684.

