## Extended Data for "DeepD3, an Open Framework for Automated Quantification of Dendritic Spines"

Extended Data Figures and Tables

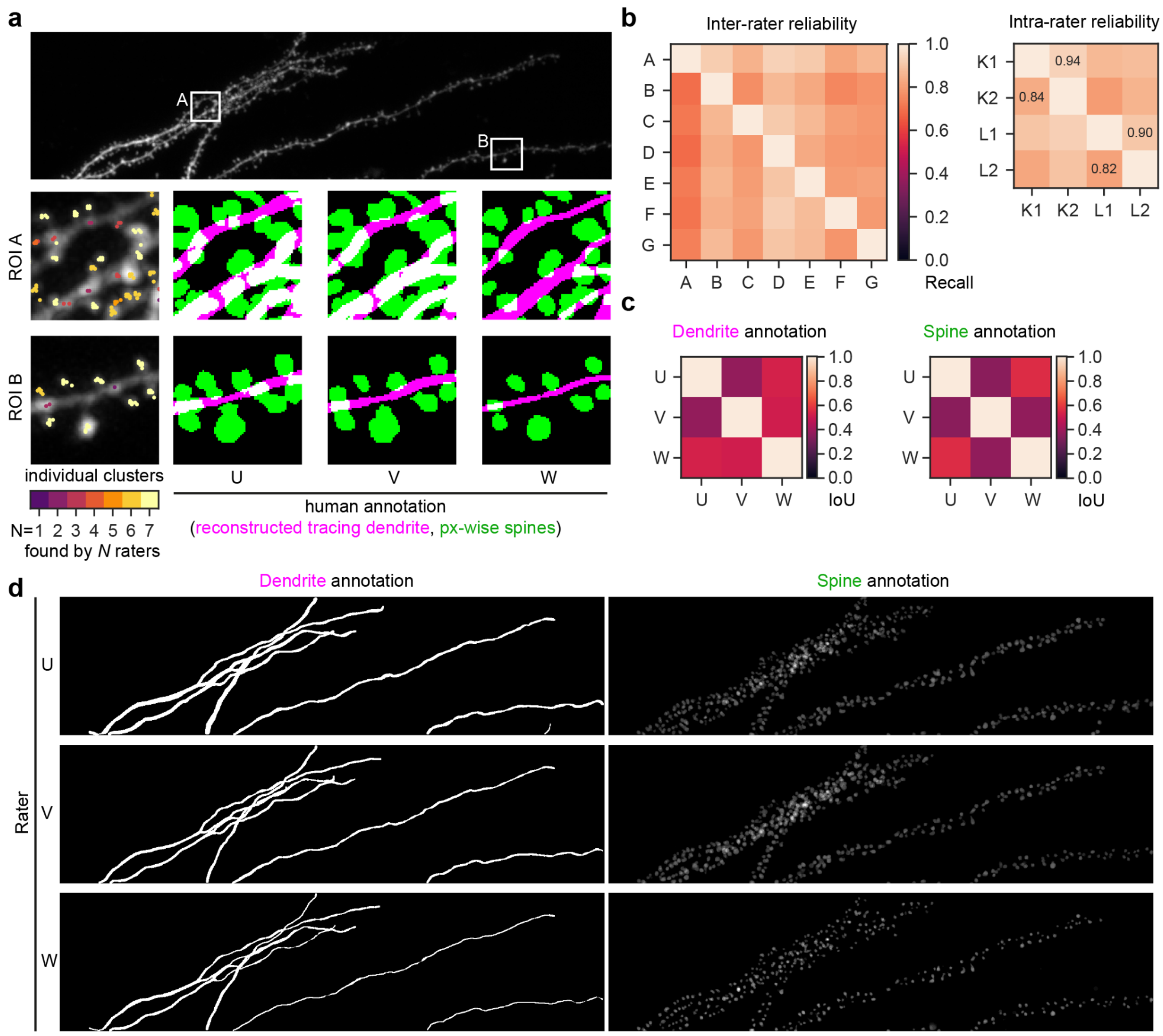


**Extended Data Figure 1. Inter-rater variability is high in spine detection, dendrite tracing and pixel-wise spine annotation.** **a**, the top panel shows a z-projection of the benchmark dataset with two regions of interest (ROIs), A and B. The two bottom panels enlarge the two ROIs and show the z-projection of the benchmark dataset stack together with single spine annotations, color-coded by the individual cluster size (left). The next three subpanels show the reconstruction of the traced dendrite (magenta) and the pixel-precise annotated dendritic spines (green) across three individual raters (U, V and W). **b**, Left, inter-rater reliability across individual raters (n=7) measured as recall. Each rater was tasked to identify a single spine by clicking on the center of mass of the spine head (see left subpanels in panel a). Right, intra-rater reliability of n=2 human experts in the benchmarking dataset. Manual annotations (1 and 2) per rater K and L were separated by at least 14 days. **c**, Intersection over union score (IoU) across individual humanannotations for reconstructed dendrite tracings (left) and pixel-precise dendritic spine annotation (right). **d**, overview of all human annotations across individual raters U, V and W.

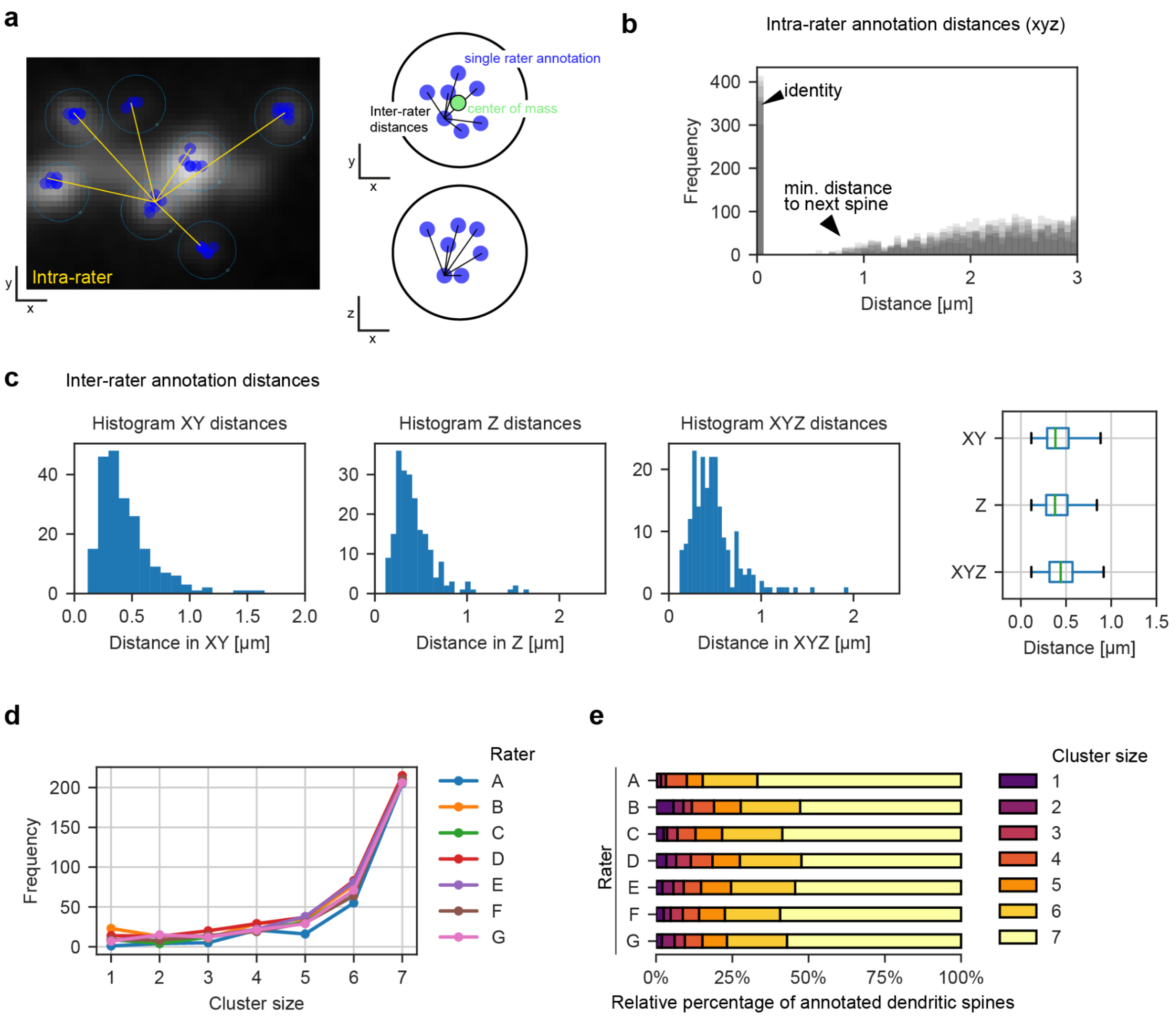


**Extended Data Figure 2. Inter- and intrarater analysis. a**, Comparison of intra- and inter-rater analyses. Left, an ROI in the benchmarking dataset with spine annotations (blue points) of 7 human experts. Distances between points of a single rater (intra-rater) are displayed in yellow. Right, distances between raters (inter-rater) are compared for a single dendritic spine in xy (top) and xz view (bottom). **b**, Three-dimensional Euclidean distance across annotated dendritic spines, overlaid across raters. Identity signifies the same dendritic spine (i.e. distance of 0). Minimal distance to the next spine starts at a Euclidean 3D distance of approximately 0.6 µm **c**, Inter-rater annotation distances for manually matched dendritic spines in µm. **d**, Frequency of found dendritic spines across raters A-G depending on the cluster size. **e**, Relative distribution of annotated dendritic spines per rater across cluster sizes. The majority of all annotated spines (> 50%) per rater are also identified by all other raters.

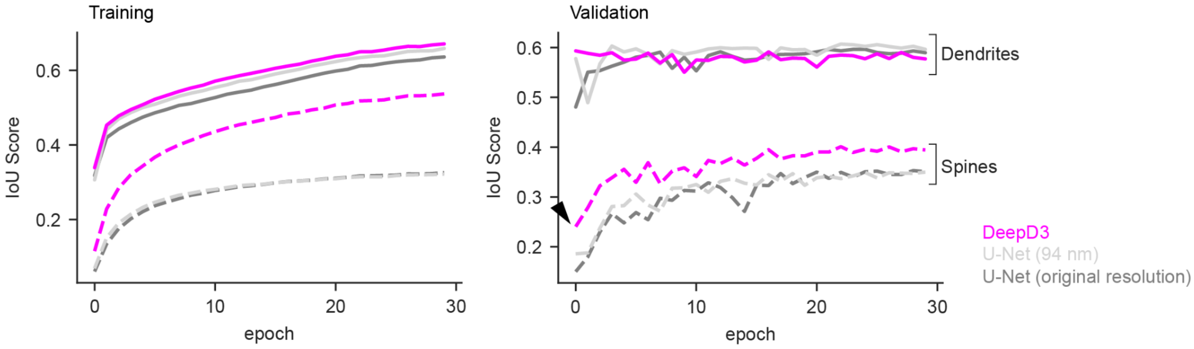


**Extended Data Figure 3. DeepD3 architecture outperforms U-Net.** The Intersection over Union (IoU) score shows the segmentation quality of the deep neural network compared to the ground truth. Performance comparison of a strong baseline (U-Net, training dataset in original resolution (dark gray) or fixed resolution (light gray)) compared to our proposed DeepD3 architecture (magenta) on the training (left) and validation (right) dataset across training epochs. NB: the DeepD3 dendritic spine performance is constantly above the U-Net baseline in the validation dataset. In addition, it converges faster and already shows superior performance after the first training epoch (black arrowhead).

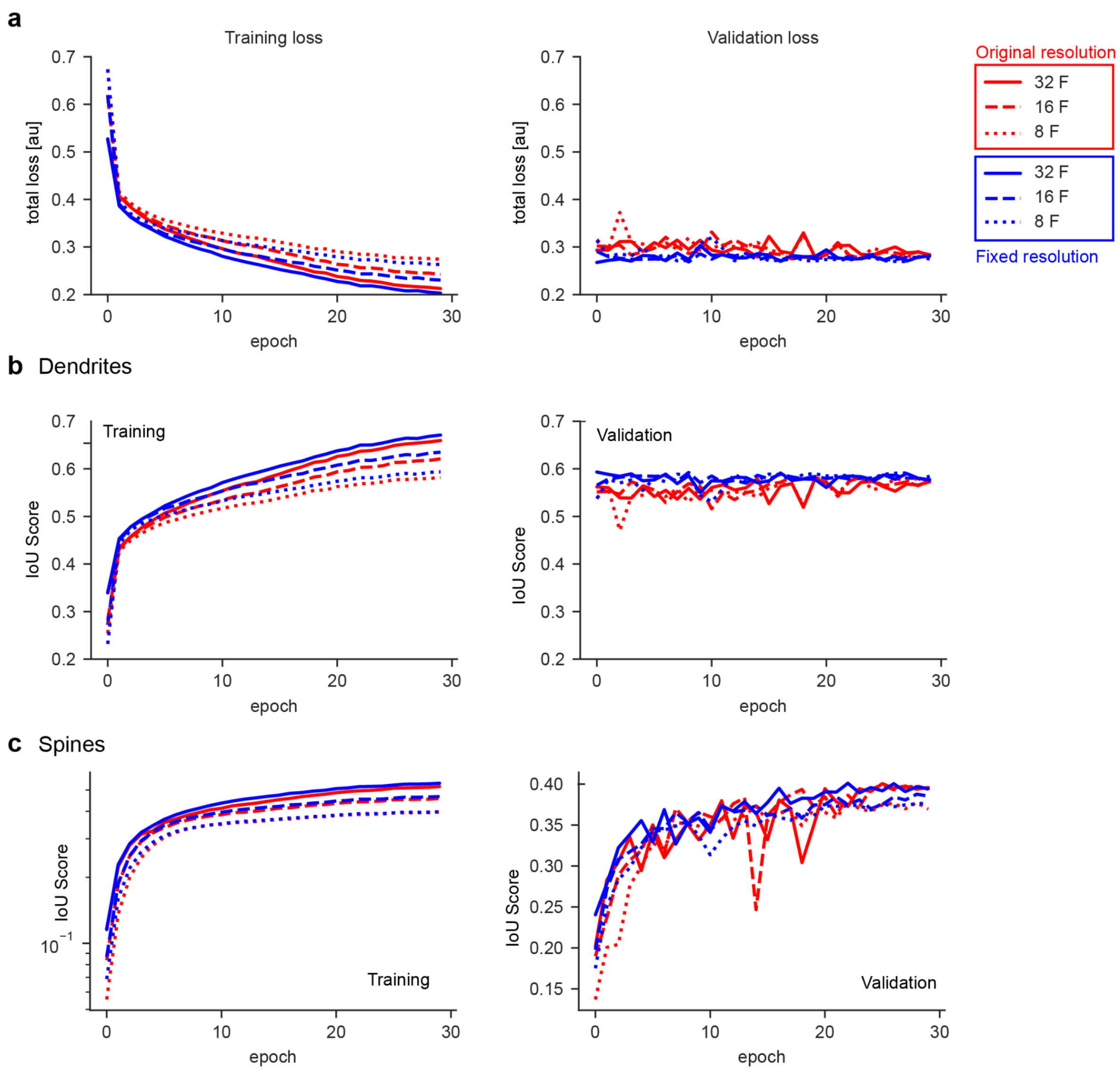


**Extended Data Figure 4. DeepD3 learns to detect dendrites and dendritic spines.**
**a**, Combined training loss for dendrites and spines (see Methods in the main manuscript). Shown are different DeepD3 scaling variants with different line styles (8F for dotted, 16 for dashed, and 32F for solid lines) for either a fixed training resolution (here 94 nm, shown in blue) or utilizing the original (mixed) training data resolutions (red). **b**, Intersection over union score (IoU) across training epochs for training data (top panel) and validation data (bottom panel). **c**, IoU score across training epochs for spines. N.B. spine IOU score converges between epochs 20-30.

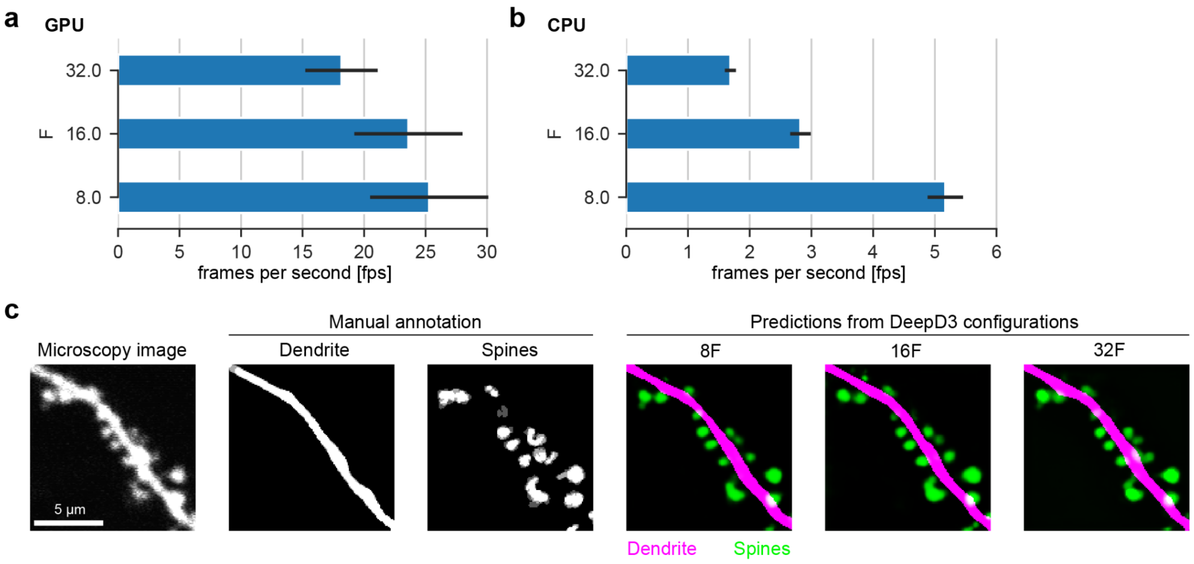


**Extended Data Figure 5. DeepD3 neural architecture allows scaling to enhance inference speed by minimally sacrificing prediction quality.**

**a**, Inference speed of a single 512x512 px tile measured in frames per second on a consumer GPU (NVIDIA RTX A4000) across DeepD3 scaling variants, denoted as 8F, 16F, and 32F. **b**, Inference speed of a single 512x512 px tile measured on a consumer CPU (AMD Ryzen 3950X) across DeepD3 scaling variants. **c**, Exemplary data tiles showing together with manual annotation and their respective network prediction (dendrite in magenta, spines in green) across DeepD3 scaling variants.

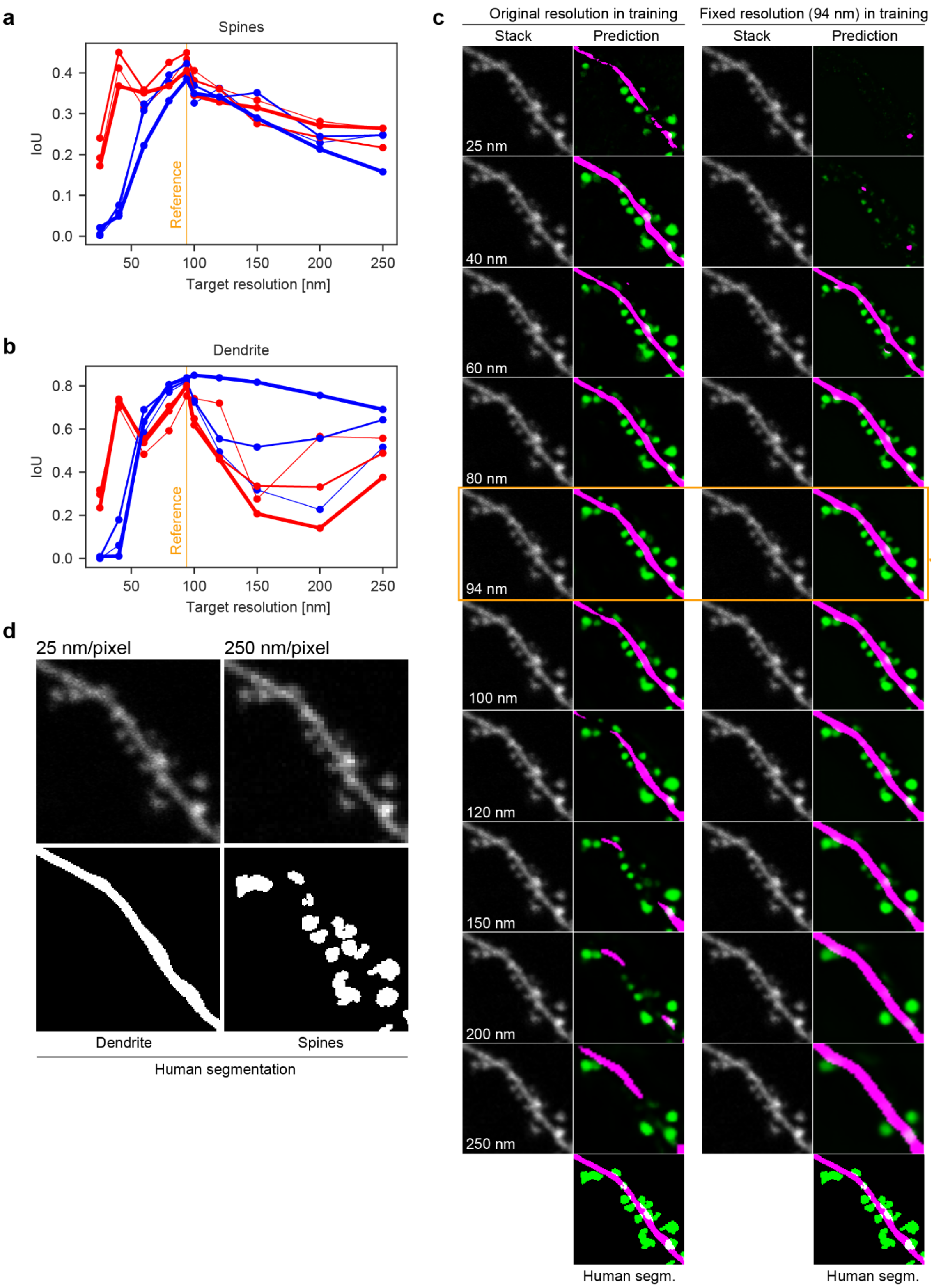


**Extended Data Figure 6. DeepD3 is stable across resolutions depending on the training paradigm.**
**a**, Intersection over Union (IoU) score for spine predictions across artificially generated resolutions from the validation dataset. Original (mixed) resolution in red, fixed resolution to 94 nm in blue. Line thickness represents used base filters (8, 16 or 32, for light, medium, or thick line strokes, respectively). **b**, Same as panel a but for dendrite predictions. **c**, Left column: Example tile of raw data of the validation dataset across resolutions. Second column DeepD3 prediction using a neural net trained on mixed resolution on raw data of various resolutions (dendrite prediction in magenta, spines prediction in green). Third column: same as left column. Fourth column: same as the second column but for a neural net trained on a fixed resolution. The bottom row shows the pixel-precise segmentation provided by a single human expert.. **d**, Two example tiles to showcase the effect of resizing a given tile to different resolutions. Top: scaled raw data. Bottom: human-segmented dendrite (left) and spine (right) segmentations of the raw data shown on top.

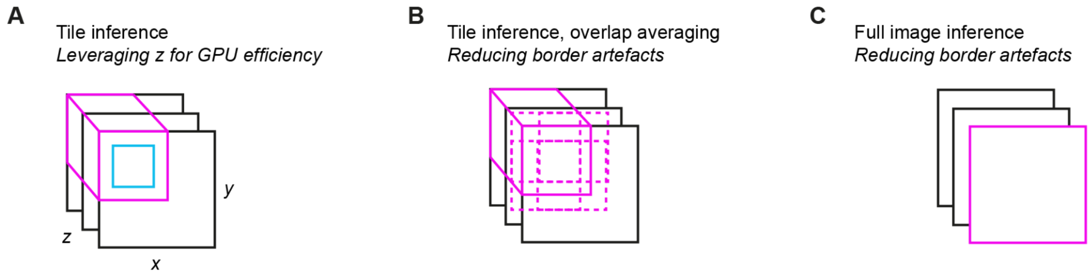


**Extended Data Figure 7. Available inference modes for the DeepD3 framework.**
**a,** Inference of image tiles (pink) that are slid in x and y. Only a smaller inset of the tile (blue rectangle) is used for the final prediction. To leverage the full GPU power, we utilize the z-depth to avoid IO-bound bottlenecks (see Online Methods). **b**, Inference of image tiles (pink) but overlapping in four directions and averaged such that border artifacts are reduced. **c**, Full image inference at a given z-depth.

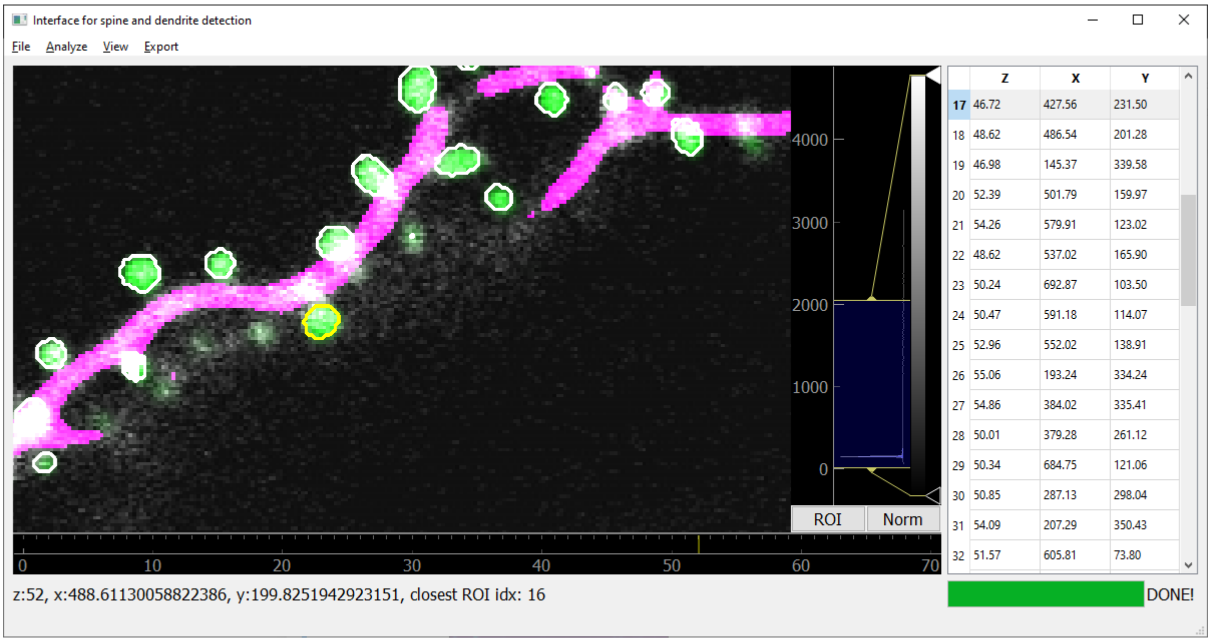


**Extended Data Figure 8. Graphical User Interface.**
Example screenshot of the DeepD3 Graphical User Interface (GUI). Tools can be selected via a drop-down menu (top). In the center part of the GUI, the loaded raw data (grayscale), performed predictions of dendritic spines (green) and dendrites (magenta), as well as generated spine ROIs (white or yellow outlines of dendritic spine predictions) are displayed. To the right of the main window, the contrast of the raw data can be adjusted (grayscale bar). Identified spine ROIs and their location in three dimensions are listed per z-level on the right side of the GUI (table with Z, X, Y columns and numbers as rows). Below is a loading bar (green) that communicates the progress of the current processing step to the user.

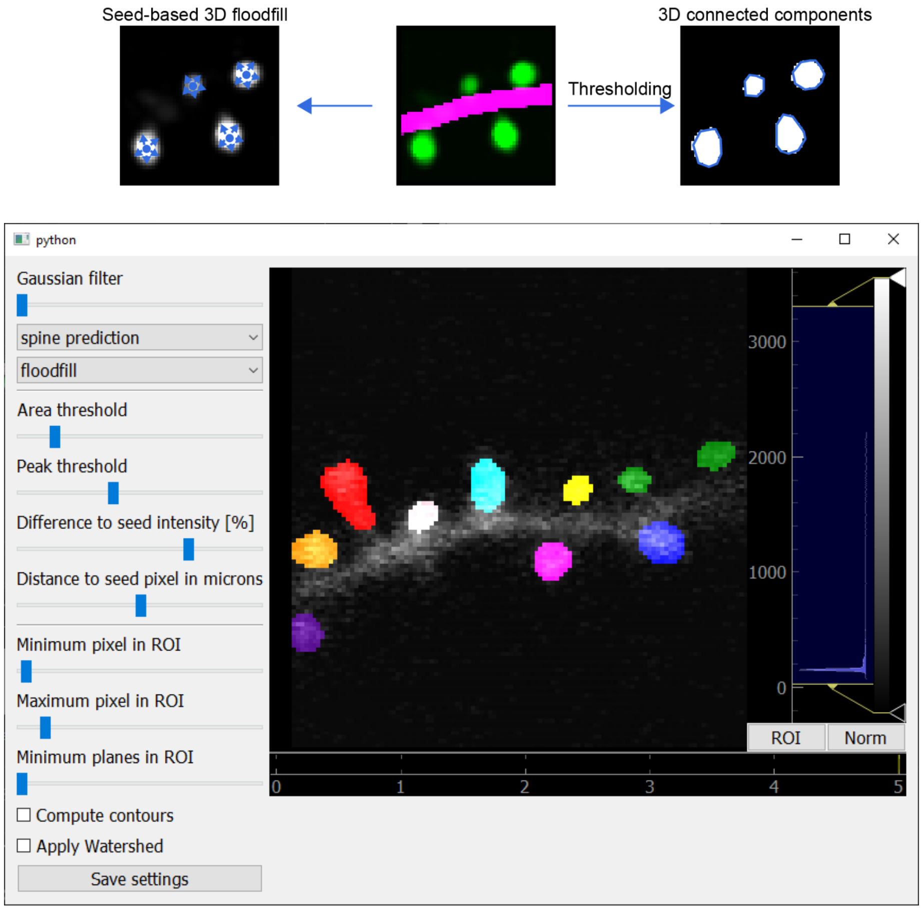


**Extended Data Figure 9. Testing 3D ROI building in Graphical User Interface.**
Top: two approaches of building spine ROIs from the cleaned spine prediction map (seed-based 3D flood-fill and 3D connected components). Bottom: DeepD3 built-in option to generate real-time feedback when building 3D ROIs to rapidly fine-tune user-defined hyperparameters. On the right side of this GUI, hyperparameters can be defined, and the main window shows the resulting spine ROIs (in different colors). Raw data are shown in grayscale and can be contrast adjusted (histogram on the right).

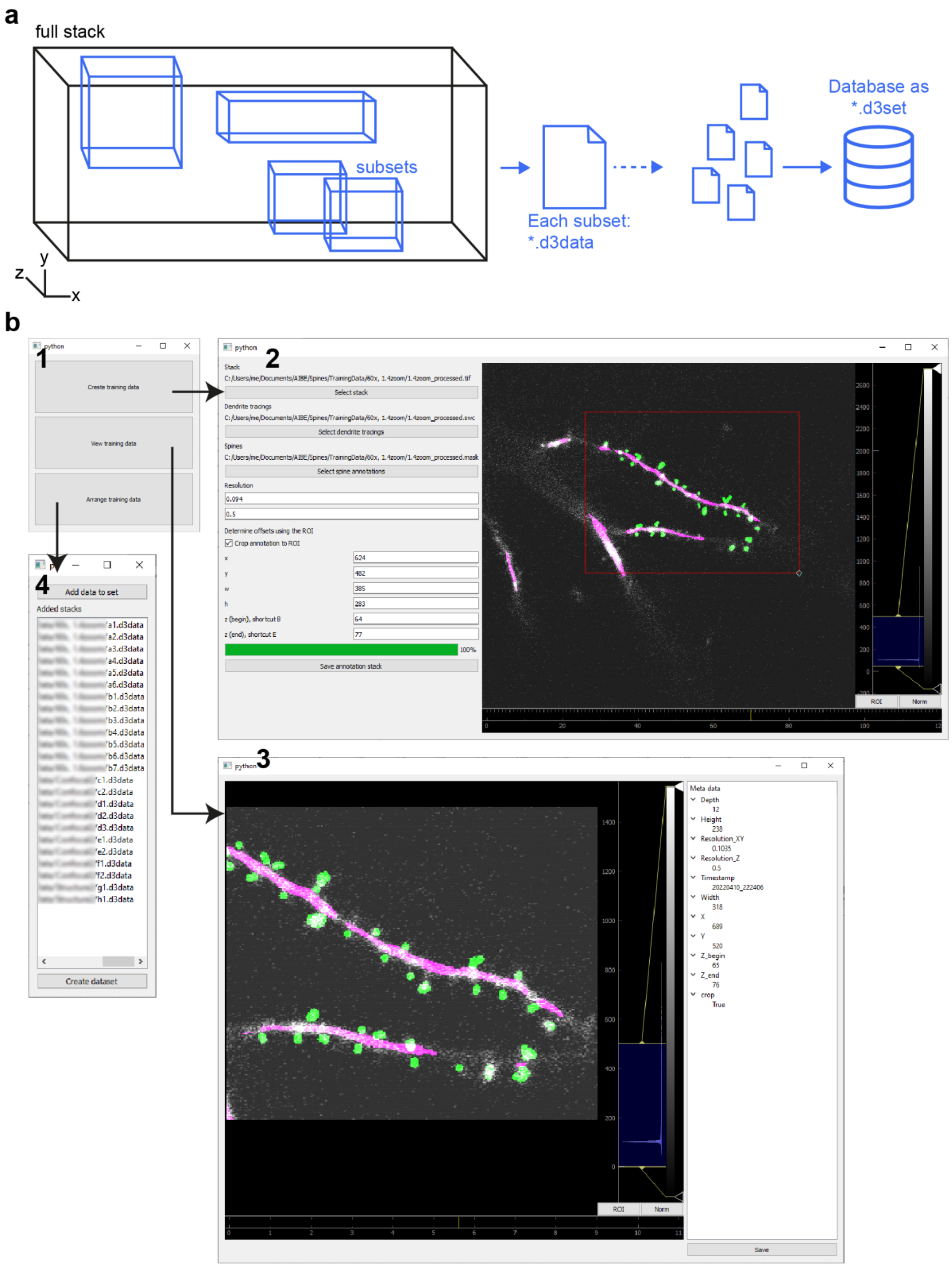


**Extended Data Figure 10. Graphical User Interface for Training Data Compilation.**
**a**, Workflow of the training data compilation pipeline. A given (partially) annotated stack (black box) can be used as training/validation/test data source. One can select one or multiple subsets (in x,y,z) each of which is saved as a *.d3data file. User-selected files can be compiled to a database-like *.d3set that contains these individual subsets. The *.d3set files can be further used in the training pipeline. **b**, Graphical User Interface elements that conveniently address the workflow in panel a. (1) is the main window, where one can access the subsequent features. (2) allows stack loading, dendrite reconstruction and subset generation, as well as *.d3data saving. (3) allows revisiting saved *.d3data subsets. (4) compiles selected files to a *.d3set.

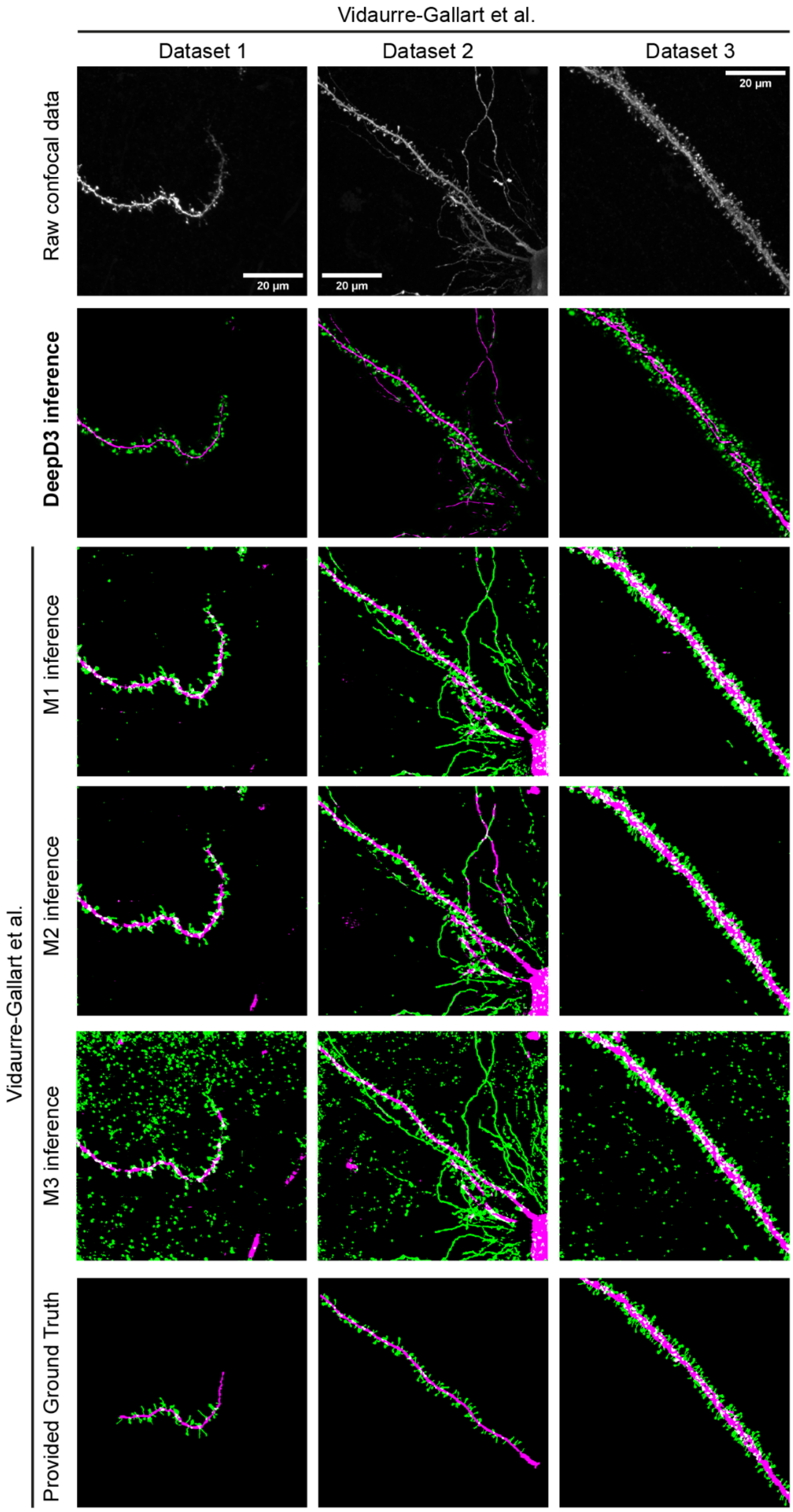


**Extended Data Figure 11. Qualitative evaluation of contemporary methodology.**
Top row: original raw TIFF stack datasets kindly provided by Vidaurre-Gallart et al. (2022). Second row: qualitative DeepD3 prediction performance on the raw data. Each plane-by-plane inference of the raw TIFF stack was performed using the DeepD3-32F network trained on unconstrained pixel resolution. Raw predictions were cleaned using the default DeepD3 cleaning settings and procedures. Third to fifth row: inference using the networks M1, M2 and M3, respectively, kindly provided by Vidaurre-Gallart et al (2022). Bottom row: ground-truth labels kindly provided by Vidaurre-Gallart et al (2022).

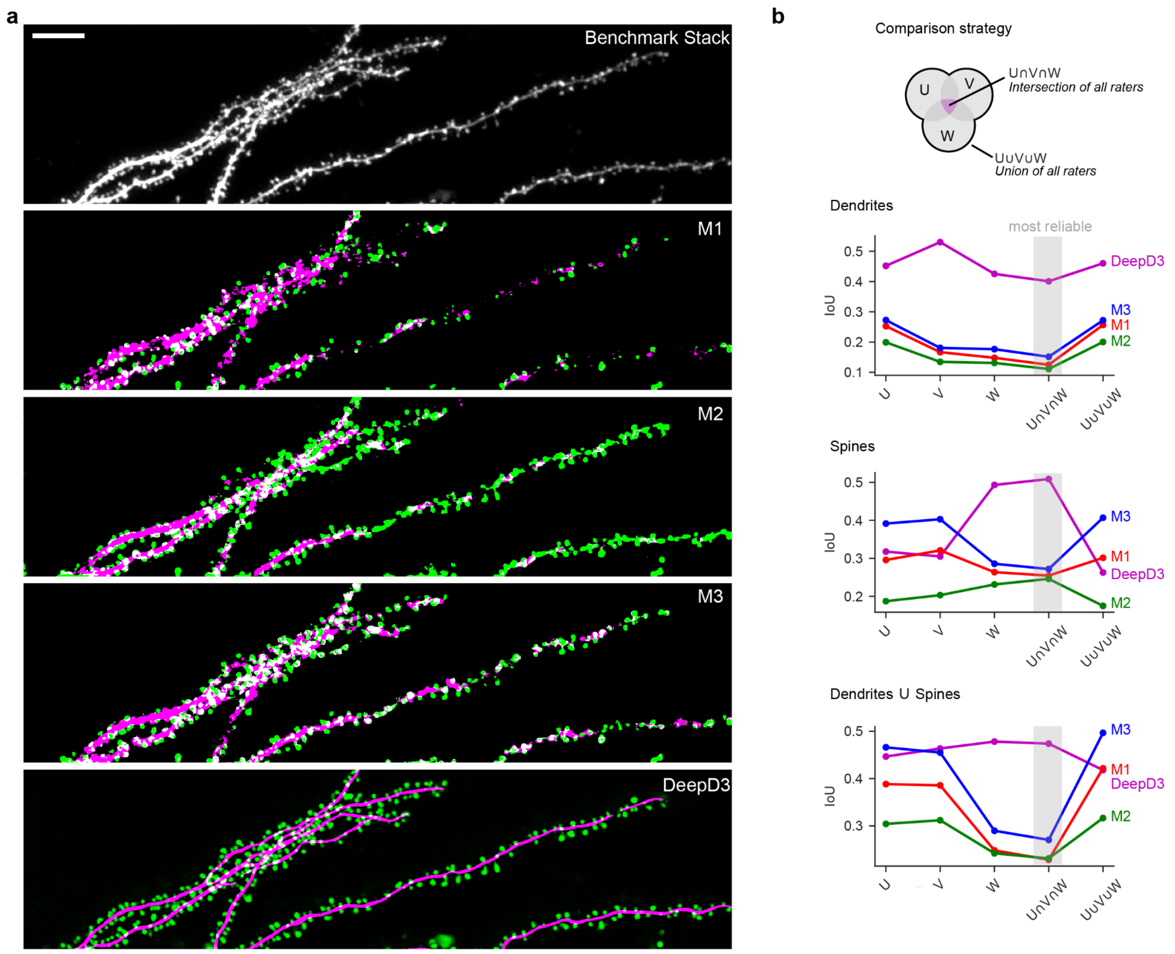


**Extended Data Figure 12. Qualitative evaluation of contemporary methodology.**
**a,** Original raw TIFF stack from the DeepD3 benchmark dataset (top) was analyzed using the methodology described in Vidaurre-Gallart et al. (2022, center three images). The DeepD3 prediction (bottom) was generated using the DeepD3-32F network trained on unconstrained pixel resolution. Raw predictions were cleaned using the default DeepD3 cleaning settings and procedures. Qualitatively, the approach presented by Vidaurre-Gallart et al., (2022) fails to accurately segment dendrites and dendritic spines in the DeepD3 benchmark dataset. Scale bar is 10 µm. **b,** Quantification of segmentation performance across individual raters, their intersection and their union annotation (see schematic on top). The IoU score is shown across neural networks M1, M2 and M3 (Vidaurre-Gallart et al., 2022) colored in red, green and blue, respectively, and DeepD3 (magenta). The intersection of all raters is thought as most reliable (indicated with shaded box). Performance is shown for spines, dendrites and the union of dendrite and spine labels.

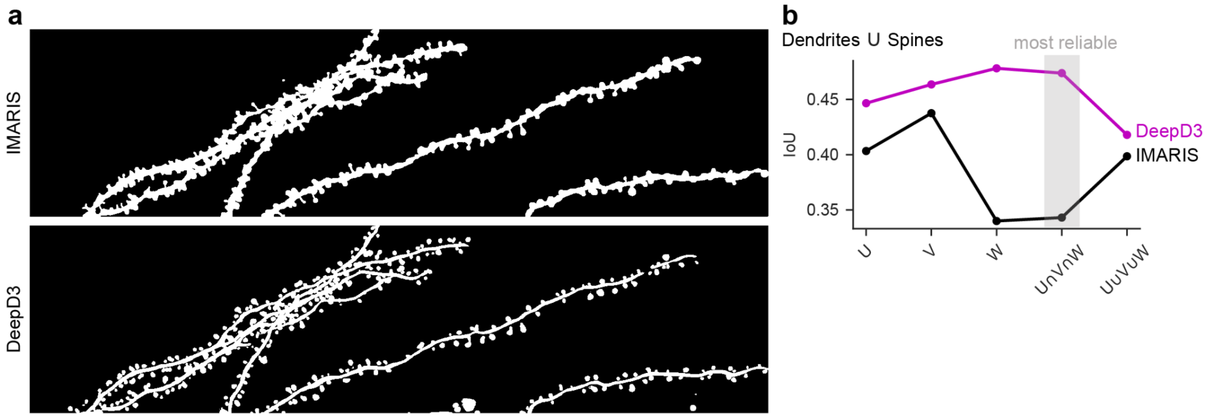


**Extended Data Figure 13: DeepD3 qualitatively and quantitatively outperforms semi-automatic state-of-the-art methods.**

**a,** Maximum intensity z-projection of the benchmark dataset with dendrites/spines extraction using IMARIS (upper panel) and DeepD3 (lower panel, union of dendrite and spine prediction with a threshold of 0.5). **b,** IoU quantification to human segmentations as introduced in Extended Data Figure 12b.

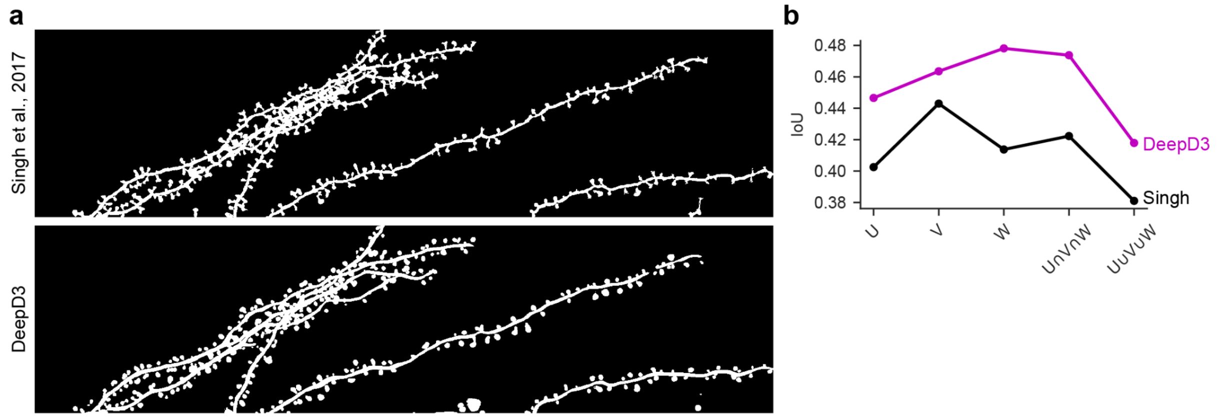


**Extended Data Figure 14: DeepD3 qualitatively and quantitatively outperforms fully automatic state-of-the-art methods.**

**a,** Maximum intensity z-projection of the benchmark dataset with dendrites/spines extraction using the method introduced by Singh et al., 2017 (upper panel) and DeepD3 (lower panel, union of dendrite and spine prediction with a threshold of 0.5). **b**, IoU quantification to human segmentations as introduced in Extended Data Figure 12b.

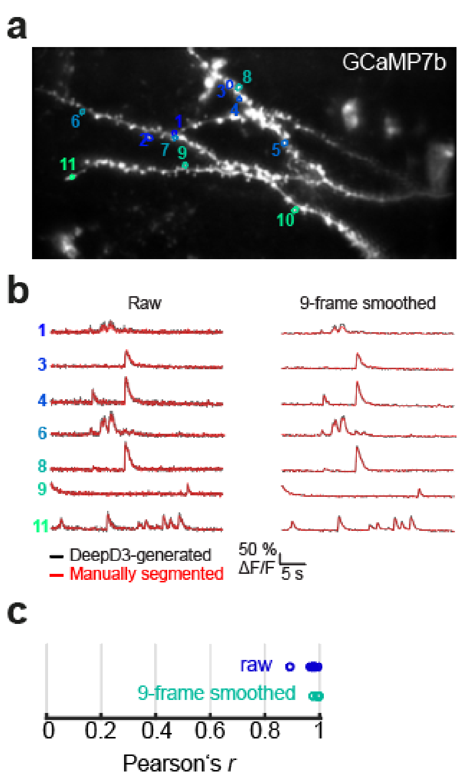


**Extended Data Figure 15. DeepD3-generated and manually segmented spine ROIs generate near-identical timecourse calcium fluctuation data.**

**a**, average projection of the analyzed calcium-imaging movie with DeepD3-generated spine ROI outlines in color and assigned numbers. **b**, Raw and 9-frame smoothed (left and right, respectively), calcium transients ($\Delta F/F_0$) of responsive spines extracted using DeepD3-generated (black) and manually segmented ROIs (red). **c**, Pearsons's correlation coefficient *r* raw (blue) and 9-frame smoothed (turquoise) traces shown in b.

| Reference | Software Name / Acronym | Brief description of spine detection | Dimensionality of input data | Notes | Code Availability |
|---|---|---|---|---|---|
| ⚫ Vidaurre-Gallart et al., 2022 | DeepSpine Tool / DeepSpine Net | 3D convolutional neural network with model options, watershed segmentation. | 3D | Second use of deep learning in spine detection. Data available upon request. Allows for manual post-editing. | Yes |
| ⚫⚫ Xiao et al., 2018 | - | Deconvolution during preprocessing, convolutional neural networks. | 2D | First use of deep learning in spine detection. | No |
| Smirnov et al., 2018 | - | Normalization, 2D median filter, Otsu thresholding to identify background, background subtraction, adaptive thresholding to binarize, skeletonization, artifact removal, skeleton smoothing, identification of disconnected spines, geodesic distance transform from dendrite to identify spine seed locations, local geodesic distance transforms to extract features, neural network-based spine identification. | 3D | Training data is open source. Does not provide full segmentation. | Yes |
| Singh et al., 2017 | - | 3D Gaussian filtering, binary thresholding of Hessian matrix, centerline extraction, seeding via voxel coding algorithm, thresholding of spine voxels. | 3D | - | Yes, but not all processing steps worked in our hands. |
| On et al., 2017 | DendritePA | Top hat filter, 2D median filter, image enhancement, segmentation using Otsu method, dendrite segmentation via piecewise convolution kernel method with low-pass filter, spine segmentation via distance map and subsequent watershed segmentation, spine categorization. | 3D + time | Approach was compared to NeuronIQ. Can also perform fluorescence-based analyses of spines (e.g. co-localization of cofilin). | Yes |
| Xie et al., 2017 | - | Image normalization, enhance linear structures, binary thresholding, skeletonization and branch point identification, spines are identified using alternative points in the skeleton. | 3D | Also detects boutons, can be utilized to identify synapses. Only performs localization, no segmentation. | Upon request |
| ⚫ Dickstein et al., 2016 | Neurolucida 360 | Based on Wearne et al., 2005 and Rodriguez et al., 2008. | 3D | - | Licensed software |
| Blumer et al., 2015 | - | Training: compute synthetic fluorescence images based on | 3D + time | Semi-automated approach. Utilized | No |

| | | | | | |
|---|---|---|---|---|---|
| | | EM data.calculate spine and dendrite probability maps based on the PCA of dendrite-orthogonal slices of synthetic images.<br><br>Method: generate 2D slices, generate spine probability maps using 9 orientation-dependent probability PCA models, binary thresholding of probability map.<br><br>Time-series analysis: rigid registration, spine matching via distance and detection probability of all possible spine paths. | | correlative light and electron microscopy (CLEM) data for benchmarking. CLEM data is no longer available. Specialized on CA1 pyramidal neurons. Requires re-training if done with a different microscope and/or cell type. | |
| Shi et al., 2014 | - | Image filtering, adaptive thresholding, morphological filtering, ray casting, wavelet transform. | 3D | Semi-automated spine classification possible. | No |
| Rada et al., 2014 | - | 2 Methods: (1) Median filter, Hessian matrix calculation, dot enhancement filter, adaptive thresholding of the original image, “morphological thinning”, skeletonization (2) generate training set using SIFT features and manual labelling, train linear SVM on manually labelled SIFT feature vectors. Spines in both methods are then segmented using watershed segmentation followed by a variational-based algorithm. | 2D | Approach was compared to NeuronIQ. | No |
| Su et al., 2014 | - | Median filter, gradient magnitude image normalization, custom Hessian filtering, white top-hat filtering to remove spines, binarization, skeletonization, dendrite boundary detection, dendrite subtraction, watershed segmentation of spines. | 2D | Requires super-resolution images as input. Approach was compared to Zhang et al., 2007 and NeuronStudio. Can perform spine density calculations. | No |
| Ortiz et al., 2014 | - | Method details unclear but based on simple linear iterative clustering resulting in supervoxels, which are then clustered using maximal inscribed spheres to segment spines. | 3D | - | No |
| Erdil et al., 2012 | - | Median filtering, edge-preserving smoothing, Otsu thresholding, extended maxima transform and boundary extraction, watershed segmentation, graph-theoretic segmentation and k-means clustering for final segmentation of spines. | 2D | - | No |

| | | | | | |
|---|---|---|---|---|---|
| He et al., 2012 | - | ‘Regularized morphological filtering’, Otsu thresholding to binarize, non-linear degeneration equation to detect spines, thresholding to segment, spine classification. | 3D | Approach was compared to Rodriguez et al., 2008, Zhang et al., 2009, and Fan et al., 2009. Can perform spine classification. | No |
| Swanger et al., 2011 | - | Semi-automated dendrite detection based on thresholds in the IMARIS software suite. | 3D + time | Time-course data analysis is possible but it is not automated; method implemented in IMARIS. | No |
| Son et al., 2011 | - | Unsharp mask filtering, binarization, anisotropic diffusion filtering followed by ISODATA algorithm to obtain skeleton of dendrites and spines, end points are detected and used as seed points for geodesic active contour modelling of spines, watershed segmentation. Spine matching across time based on Lucas-Kanade method (optical flow of spine within predetermined region of another spine at the previous time point). | 2D + time | Approach was compared to NeuronIQ. Offers automatic spine classification. | No |
| Mukai et al., 2011 | Spiso-3D | Deconvolution, scale-space transformation, dendrite ridge line detection and subsequent 3D dendrite reconstruction, 3D dendrite subtraction and subsequent spine point determination via eigenvalues of the Hessian tensor, spine diameter calculation, manual correction. | 3D | Semi-automated approach. Only performs localization, spine neck length and spine diameter calculations, no full segmentation. | No |
| Zhang et al., 2010 | - | generalized gradient vector flow, eigen-analysis, multiple shape measurements and fast marching algorithm. | 3D | - | No |
| Zhang et al., 2010 | - | Iterative deconvolution, Otsu thresholding, gradient vector field calculation, feature point detection, spine detection via eigen-analysis method, segmentation via fast marching method, postprocessing. | 3D | Intended to be used on medium-sized spiny neurons. | No |
| Li et al., 2010 | - | Spine detection: Hessian matrix, curvilinear structure detection to identify dendrite, length-based criterion for spines and dendrites<br><br>Registration: iterative closest reciprocal point algorithm-based rigid transformation followed by non-rigid local deformation for | 2D + time | This approach focuses on chronic data quantification. Approach was compared to Li et al., 2009, Koh et al., 2002, and Fan et al., 2009. First time an approach deals | No |

| | | | | | |
|---|---|---|---|---|---|
| | | fine-scale registration.<br><br>Spine matching: distance-based criterion, exclusive matching between time points, global similarity metric. | | with dis- and re-appearing spines. Dataset available upon request. | |
| Fan et al., 2009 | - | Curvilinear dendrite detection, adaptive local binary fitting Level Set Model, Laplacian of Gaussian, Maximum Likelihood Estimation. | 3D + time | Time-course data analysis possible. | Upon request |
| ⚫<br>Yuan et al., 2009 | Part of the FARSIGHT toolkit | Deconvolution, grayscale skeletonization, high-curvature and critical point detection, iterative path-line formation algorithm, Graph Generation & Computation of the Intensity-weighted Minimal Spanning Tree (IW-IMST), MDL-Based Estimation of Dendritic Backbones and spines. | 3D | Evaluated on data from multiple laboratories. Informative section on related literature. | Available upon request, available open-source FARSIGHT toolkit |
| Li et al., 2009 | - | 3D median filter, top-hat filter, Rayburst sampling using user-defined seed points, fuzzy C-mean clustering, iterative low-pass filtering and mesh decimation, spine detection via region growing followed by watershed segmentation. | 3D | Semi-automated. Approach was compared to NeuronStudio. Claims to be suitable for images containing multiple neurons/dendrites. | No |
| Janoos et al., 2009 | - | Deconvolution, non-linear diffusion filtering, binarization, set distance boundaries and reconnect floating spine heads using active contour shape models, extended marching cubes algorithm to surface dendrites and spines, skeleton extraction using a medial geodesic function, spine detection using length-based branching criterion on skeleton. | 3D | - | No |
| ⚫⚫<br>Rodriguez et al., 2008 | NeuronStudio | Adaptive local thresholding, voxel clustering and Rayburst Sampling. | 3D | - | No longer available |
| Zhang et al., 2007 | - | Curvilinear structure detection, linear discriminate analysis. | 2D | - | No |
| ⚫<br>Cheng et al., 2007 | NeuronIQ | adaptive thresholding, backbone extraction, local-region-cutting algorithm, SNR-based threshold. | 3D | - | No longer available |
| Bai et al., 2007 | - | Deconvolution, unsharp mask filtering, 3D median filtering, binarization, size-dependent connected component analysis to determine dendrites and spines, skeletonization, angle- and width-based criterion on skeleton to identify and segment dendritic | 2D | Can also compute spine length. | No longer available |

| | | | | | |
|---|---|---|---|---|---|
| | | spines. | | | |
| ⚫⚫ Xu et al., 2006 | NeuronIQ | Median filter, deconvolution, unsharp masking and subsequent binarization, medial axis transform to identify the skeleton, angle- and length-based exclusion of spine segments of the skeleton, grassfire algorithm to detect potential spine seed points, second grassfire algorithm to detect dendrite boundary. | 3D | - | No |
| ⚫⚫ Wearne et al., 2005 | Part of the NeuronStudio package | Deconvolution, thresholding, skeletonization and subsequent medial axis calculation, Rayburst sampling, iterative thinning algorithm, custom cleanup of dendritic tree skeleton. | 3D | Primarily utilized for neural morphology analysis. Basis for Rodriguez et al., 2008. | Licensed software |
| ⚫ Weaver et al., 2004 | 3DMA-neuron | Deconvolution, 2x voxel compression in z, semi-automated data registration, manual tiling, binarization, skeletonization, dendrite radius estimation, spine detection using protrusions of spines from the dendrite as potential seed points, manual revision option at end. | 3D + time | Further refined Koh et al., 2002 to extend use cases to large datasets (e.g. entire neuron data). | No longer available |
| ⚫⚫ Koh et al., 2002 | 3DMA-neuron | Deconvolution, dendritic backbone extraction via medial axis algorithm. Active contour model, volume-based matching, spine categorization. | 3D + time | Relatively slow due to computational burden. | No longer available |
| ⚫⚫ Koh and Lindquist, 2001 | 3DMA-neuron | Deconvolution, dendritic backbone extraction via medial axis algorithm. Active contour model, volume-based matching, spine categorization. | 3D + time | Relatively slow due to computational burden. | No longer available |
| ⚫⚫ Watzel et al., 1995 | - | Binarization, skeletonization, dendrite reconstruction and subsequent subtraction from binary image, generation of spine seeds by finding skeletal points adjacent to dendrite, seed classification, minimal box placement around each spine. | 3D | One of the first efforts to automate spine detection. Only works with one unbranched dendrite with all spines of interest connected to the dendrite. | No |
| ⚫ Rusakov and Stewart, 1995 | - | Region of interest selection, binarization via valley global minimum detection, skeletonization, break bifurcation points and measure segments, manual classification of segments into spine, dendrite and “ignored”. | 2D | One of the first efforts to automate spine detection. Semi-automated approach. Can perform spine density estimation. | No |

**Extended Data Table 1. Previous approaches of (semi)-automated spine detection.**
⚫⚫ indicate landmark papers in the field. ⚫ denote significant contributions to the field.

| Dataset | Species | Mode | Acquisition technique | Dye/ Fluorophore | XY resolution [μm] | Z step [μm] |
|---|---|---|---|---|---|---|
| DeepD3 Benchmark | Rat | Ex vivo | Two-photon | tdTomato | 0.094 | 0.50 |
| External A | Mouse | In vivo | Two-photon | iGluSnFR | 0.200 | 0.75 |
| External B | Mouse | In vivo | Two-photon | Thy1-YFP | 0.117 | 0.50 |
| External C | Human | In vitro | Confocal | Biocytin + HRP + DAB | 0.240 | 0.42 |

**Extended Data Table 2. Datasets that were evaluated in this study.** Data of External A was described in Kazemipour et al., 2019. Data of External B was described in Frank et al., 2018. Data of External C was described in Peng et al., 2015 and Manubens-Gil et al., 2022. HRP = Horseradish peroxidase, DAB = 3,3'-Diaminobenyidine tetrahydrochloride.

| Model | $f_{base}$ | Trained on | Output |
|---|---|---|---|
| DeepD3_8F_94nm.h5 | 8 | Complete training data, but rescaled to match an xy-resolution of 94 nm | Spine and dendrite prediction maps |
| DeepD3_16F_94nm.h5 | 16 | | |
| DeepD3_32F_94nm.h5 | 32 | | |
| DeepD3_8F.h5 | 8 | Complete training data with various xy-resolutions | |
| DeepD3_16F.h5 | 16 | | |
| DeepD3_32F.h5 | 32 | | |

**Extended Data Table 3. Model Zoo.** All models were trained on the DeepD3 training dataset using the different $f_{base}$ configurations. We offer two sets of models: trained with original resolution and trained on rescaled data to match a 94 nm resolution. Each network yield two prediction maps: dendrites and dendritic spines with a probability between 0 (px does not belong to class) and 1 (px belongs certainly to class).

### Extended Data References


Bai, W., Zhou, X., Ji, L., Cheng, J., & Wong, S. T. (2007). Automatic dendritic spine analysis in two-photon laser scanning microscopy images. *Cytometry Part A: The Journal of the International Society for Analytical Cytology*, *71*(10), 818–826.

Blumer, C., Vivien, C., Genoud, C., Perez-Alvarez, A., Wiegert, J. S., Vetter, T., & Oertner, T. G. (2015). Automated analysis of spine dynamics on live ca1 pyramidal cells. *Medical image analysis*, *19*(1), 87–97.

Cheng, J., Zhou, X., Miller, E., Witt, R. M., Zhu, J., Sabatini, B. L., & Wong, S. T. (2007). A novel computational approach for automatic dendrite spines detection in two-photon laser scan microscopy. *Journal of neuroscience methods*, *165*(1), 122–134.

Dickstein, D. L., Dickstein, D. R., Janssen, W. G., Hof, P. R., Glaser, J. R., Rodriguez, A., O'Connor, N., Angstman, P., & Tappan, S. J. (2016). Automatic dendritic spine quantification from confocal data with neurolucida 360. *Current protocols in neuroscience*, *77*(1), 1–27.

Erdil, E., Yagci, A. M., Argunsah, A. O., Ramiro-Cort´es, Y., Hobbiss, A. F.,¨ Israely, I., & Unay, D. (2012). A tool for automatic dendritic spine detection and analysis. part i: Dendritic spine detection using multilevel region-based segmentation. *2012 3rd International Conference on Image Processing Theory, Tools and Applications (IPTA)*, 167–171.

Fan, J., Zhou, X., Dy, J. G., Zhang, Y., & Wong, S. T. (2009). An automated pipeline for dendrite spine detection and tracking of 3d optical microscopy neuron images of in vivo mouse models. *Neuroinformatics*, *7*(2), 113–130.

Frank, A. C., Huang, S., Zhou, M., Gdalyahu, A., Kastellakis, G., Silva, T. K., Lu, E., Wen, X., Poirazi, P., Trachtenberg, J. T., et al. (2018). Hotspots of dendritic spine turnover facilitate clustered spine addition and learning and memory. *Nature communications*, *9*(1), 422.

He, T., Xue, Z., Kim, Y., & Wong, S. T. (2012). Three-dimensional dendritic spine detection based on minimal cross-sectional curvature. *2012 9th IEEE International Symposium on Biomedical Imaging (ISBI)*, 1639–1642.

He, T., Xue, Z., & Wong, S. T. (2012). A novel approach for three dimensional dendrite spine segmentation and classification. *Medical Imaging 2012: Image Processing*, *8314*, 919–926.

Janoos, F., Mosaliganti, K., Xu, X., Machiraju, R., Huang, K., & Wong, S. T. (2009). Robust 3d reconstruction and identification of dendritic spines from optical microscopy imaging. *Medical image analysis*, *13*(1), 167–179.

Kazemipour, A., Novak, O., Flickinger, D., Marvin, J. S., Abdelfattah, A. S., King, J., Borden, P. M., Kim, J. J., Al-Abdullatif, S. H., Deal, P. E., et al. (2019). Kilohertz frame-rate two-photon tomography. *Nature methods*, *16*(8), 778–786.

Koh, I. Y., & Lindquist, W. B. (2001). Automated 3d dendritic spine detection and analysis from two-photon microscopy. *Three-Dimensional and Multidimensional Microscopy: Image Acquisition and Processing VIII*, *4261*, 48–59.

Koh, I. Y., Lindquist, W. B., Zito, K., Nimchinsky, E. A., & Svoboda, K. (2002). An image analysis algorithm for dendritic spines. *Neural computation*, *14*(6), 1283–1310.

Li, K., Miller, E. D., Chen, M., Kanade, T., Weiss, L. E., & Campbell, P. G. (2008). Cell population tracking and lineage construction with spatiotemporal context. *Medical image analysis*, *12*(5), 546–566.

Li, Q., Deng, Z., Zhang, Y., Zhou, X., Nagerl, U. V., & Wong, S. T. (2010). A global spatial similarity optimization scheme to track large numbers of dendritic spines in time-lapse confocal microscopy. *IEEE Transactions on Medical Imaging*, *30*(3), 632–641.

Li, Q., Zhou, X., Deng, Z., Baron, M., Teylan, M. A., Kim, Y., & Wong, S. T. (2009). A novel surface-based geometric approach for 3d dendritic spine detection from multi-photon excitation microscopy images. *2009 IEEE International Symposium on Biomedical Imaging: From Nano to Macro*, 1255–1258.

Manubens-Gil, L., Zhou, Z., Chen, H., Ramanathan, A., Liu, X., Liu, Y., Bria, A., Gillette, T., Ruan, Z., Yang, J., et al. (2022). Bigneuron: A resource to benchmark and predict best-performing algorithms for automated reconstruction of neuronal morphology. *bioRxiv*.

Mukai, H., Hatanaka, Y., Mitsuhashi, K., Hojo, Y., Komatsuzaki, Y., Sato, R., Murakami, G., Kimoto, T., & Kawato, S. (2011). Automated analysis of spines from confocal laser microscopy images: Application to the discrimination of androgen and estrogen effects on spinogenesis. *Cerebral cortex*, *21*(12), 2704–2711.

On, V., Zahedi, A., Ethell, I. M., & Bhanu, B. (2017). Automated spatiotemporal analysis of dendritic spines and related protein dynamics. *Plos one*, *12*(8), e0182958.

Ortiz, C. A., Gonzalo-Martı, C., Pen˜a, J. M., & Menasalvas, E. (2014). 3d dendrite spine detection-a supervoxel based approach. In *Rough sets and intelligent systems paradigms* (pp. 359–366). Springer.

Peng, H., Hawrylycz, M., Roskams, J., Hill, S., Spruston, N., Meijering, E., & Ascoli, G. A. (2015). Bigneuron: Large-scale 3d neuron reconstruction from optical microscopy images. *Neuron*, *87*(2), 252–256.

Rada, L., Erdil, E., Argunsah, A. O., Unay, D., & Cetin, M. (2014). Automatic dendritic spine detection using multiscale dot enhancement filters and sift features. *2014 IEEE International Conference on Image Processing (ICIP)*, 26–30.

Rodriguez, A., Ehlenberger, D. B., Dickstein, D. L., Hof, P. R., & Wearne, S. L. (2008). Automated three-dimensional detection and shape classification of dendritic spines from fluorescence microscopy images. *PloS one*, *3*(4), e1997.

Rusakov, D. A., & Stewart, M. G. (1995). Quantification of dendritic spine populations using image analysis and a tilting disector. *Journal of neuroscience methods*, *60*(1-2), 11–21.

Shi, P., Huang, Y., & Hong, J. (2014). Automated three-dimensional reconstruction and morphological analysis of dendritic spines based on semisupervised learning. *Biomedical optics express*, *5*(5), 1541–1553.

Singh, P. K., Hernandez-Herrera, P., Labate, D., & Papadakis, M. (2017). Automated 3-d detection of dendritic spines from in vivo two-photon image stacks. *Neuroinformatics*, *15*(4), 303–319.

Smirnov, M. S., Garrett, T. R., & Yasuda, R. (2018). An open-source tool for analysis and automatic identification of dendritic spines using machine learning. *Plos one*, *13*(7), e0199589.

Su, R., Sun, C., Zhang, C., & Pham, T. D. (2014). A novel method for dendritic spines detection based on directional morphological filter and shortest path. *Computerized Medical Imaging and Graphics*, *38*(8), 793–802.

Swanger, S. A., Yao, X., Gross, C., & Bassell, G. J. (2011). Automated 4d analysis of dendritic spine morphology: Applications to stimulus-induced spine remodeling and pharmacological rescue in a disease model. *Molecular brain*, *4*(1), 1–14.

Vidaurre-Gallart, I., Fernaud-Espinosa, I., Cosmin-Toader, N., Talavera-Martınez, L., Martin-Abadal, M., Benavides-Piccione, R., Gonzalez-Cid, Y., Pastor, L., DeFelipe, J., & Garcıa-Lorenzo, M. (2022). A deep learning-based workflow for dendritic spine segmentation. *Frontiers in neuroanatomy*, *16*, 817903–817903.

Watzel, R., Braun, K., Hess, A., Scheich, H., & Zuschratter, W. (1995). Detection of dendritic spines in 3-dimensional images. In *Mustererkennung 1995* (pp. 160–167). Springer.

Wearne, S., Rodriguez, A., Ehlenberger, D., Rocher, A., Henderson, S., & Hof, P. (2005). New techniques for imaging, digitization and analysis of three-dimensional neural morphology on multiple scales. *Neuroscience*, *136*(3), 661–680.

Weaver, C. M., Hof, P. R., Wearne, S. L., & Lindquist, W. B. (2004). Automated algorithms for multiscale morphometry of neuronal dendrites. *Neural computation*, *16*(7), 1353–1383.

Xiao, X., Djurisic, M., Hoogi, A., Sapp, R. W., Shatz, C. J., & Rubin, D. L. (2018). Automated dendritic spine detection using convolutional neural networks on maximum intensity projected microscopic volumes. *Journal of neuroscience methods*, *309*, 25–34.

Xie, Q., Chen, X., Deng, H., Liu, D., Sun, Y., Zhou, X., Yang, Y., & Han, H. (2017). An automated pipeline for bouton, spine, and synapse detection of in vivo two-photon images. *BioData Mining*, *10*(1), 1–23.

Xu, X., Cheng, J., Witt, R. M., Sabatini, B. L., & Wong, S. T. (2006). A shape analysis method to detect dendritic spine in 3d optical microscopy image. *3rd IEEE International Symposium on Biomedical Imaging: Nano to Macro, 2006.*, 554–557.

Yuan, X., Trachtenberg, J. T., Potter, S. M., & Roysam, B. (2009). Mdl constrained 3-d grayscale skeletonization algorithm for automated extraction of dendrites and spines from fluorescence confocal images. *Neuroinformatics*, *7*(4), 213–232.

Zhang, Y., Chen, K., Baron, M., Teylan, M. A., Kim, Y., Song, Z., Greengard, P., & Wong, S. T. (2010). A neurocomputational method for fully automated 3d dendritic spine detection and segmentation of medium-sized spiny neurons. *Neuroimage*, *50*(4), 1472–1484.

Zhang, Y., Zhou, X., Witt, R. M., Sabatini, B. L., Adjeroh, D., & Wong, S. T. (2007). Dendritic spine detection using curvilinear structure detector and lda classifier. *Neuroimage*, *36*(2), 346–360.
