## Supplementary Note for "DeepD3, an Open Framework for Automated Quantification of Dendritic Spines"

### SUPPLEMENTARY NOTES

#### DeepD3 design and workflow

DeepD3 is an open deep-learning-based framework for the automated detection of dendritic spines and dendrites in 2D or 3D microscopy data. The full DeepD3 workflow is described here and is implemented in the graphical user interface (GUI) and the DeepD3 batch processing function (see below). After loading the image data in the GUI, a network model of choice is selected to infer the absence or presence of dendritic spines and dendrites in that image.

#### Network models

We provide variants of the DeepD3 architecture by scaling the convolutional filters based on the first layer filters (see Methods, and Extended Data Figure 4 and 5), which allow speed/quality trade offs. Lean deep neural networks with lower parameter space perform slightly worse during training and validation (Extended Data Figure 4), however, do show qualitatively similar results (Extended Data Figure 4b). The full DeepD3 architecture (32F) pays more attention to fine details (Extended Data Figure 5c), whereas the smaller architecture variants (16F and 8F) have faster inference times on both, CPUs and GPUs (Extended Data Figure 5a,b). In general, we observe an up to 1.39x (GPU) and 3.06x speed-up (CPU) when using the smaller variants (Extended Data Figure 5a, b).

#### Image inference and generation of raw prediction maps

DeepD3 is shipped with three different inference modes (Extended Data Figure 7). The user is able to select between a tile inference mode (Extended Data Figure 7a), which can be extended by averaging overlapping regions (Extended Data Figure 7b), as proposed by Ronneberger et al. (2015), and a full plane inference mode (Extended Data Figure 7c). The latter is set as default, as this reduces border artifacts. For network inference, we treat each plane independently as a two-dimensional image. Three-dimensional segmentation techniques are error-prone and tend to heavily overfit, also on dendritic spine segmentation (Vidaurre-Gallart et al., 2022), and need specific z-steps that constrain experimental environments. We show that our method nevertheless provides qualitatively high prediction accuracy throughout 3D stacks (Supplementary Movie 1) as shown exemplarily in Figure 2a and Extended Data Figure 5-6. The pre-trained networks can all be employed using the DeepD3 graphical user interface (Extended Data Figure 5), a command line tool, or as Python library, together with two 2D and 3D ROI building tools (Extended Data Figure 8).

Regardless of the type of inference mode chosen, DeepD3 inference generates two raw prediction maps for a given input image, one each for dendritic spines and dendrites. These prediction maps contain the likelihood of each pixel being part of a dendrite or dendritic spine, respectively (values between 0 and 1). The DeepD3 GUI offers flexible export options for these raw prediction maps. In the next steps, the raw prediction maps are cleaned to identify dendrites and dendritic spine ROIs using a number of user-defined hyperparameters.

#### Cleaning of the dendrite prediction map

First, the dendrite threshold is defined, which is utilized to binarize the dendrite prediction map. Values above the dendrite threshold are considered for further analysis, while values below are not. Next, the minimum dendrite size is defined (in number of pixels) to exclude those elements in the dendrite prediction map that are too small to be considered a dendrite. This is implemented via 2D/3D connected component analysis and comparing the size of the resulting components to the minimum dendrite size. We further incorporate the possibility to connect dendritic elements that have only a few pixels offset to each other to enhance the segmentation quality. Once the dendrite prediction map is cleaned, it is utilized to further refine the spine prediction map. It is of note that dendrite cleaning is optional but highly recommended as spurious dendrite prediction can negatively influence spine prediction.

#### Cleaning of the spine prediction map

Since dendritic spines grow out from a parent dendrite, typically, spines are in close spatial proximity to dendrites in most image data. This is leveraged to detect potentially erroneously predicted dendritic spines and remove them during cleaning of the spine prediction maps. To this end, a user-defined dendrite proximity threshold is utilized to dilate the cleaned dendritic prediction map. Using this dilated map as a mask for the spine prediction map generates a cleaned spine prediction map, in which all spine predictions that are outside of the dendrite proximity threshold are excluded. Hence, the dendrite proximity threshold acts as a distance-to-dendrite metric of each spine prediction, allowing the user to exclude spine predictions that are not in any spatial proximity to a detected dendrite.

#### ROI detection

The cleaned spine prediction map is utilized to generate regions of interest (ROIs) of individual dendritic spines in either 2D or 3D using either a flood-filling approach or connected components (Extended Data Figure 9). Both methods allow high-quality spine segmentations. However, we recommend using our testing environment inside of the DeepD3 GUI to experiment with hyperparameters and the ROI building methods (Extended Data Figure 9).

The flood-filling method is set up as follows: initially, all pixels that are below the area threshold (a user-set probability threshold) are excluded from the post-processed spine prediction map. Next, seed pixels are selected based on the peak threshold (a user-set threshold to determine seed pixels of novel ROIs). Seeding of ROIs is performed via a flood-filling algorithm that mines a 26-neighbor vicinity starting with the highest probability seed pixel. We iteratively assign pixels in 3D to a given ROI that fulfill the following criteria: (1) spine prediction above user-set threshold, (2) maximum euclidean distance to seed pixel, (3) spine probability relative to seed pixel's spine probability. When using the connected components approach, the spine prediction is binarized using a user-set threshold. 3D connected components are build using the cc3d library (https://pypi.org/project/connected-components-3d/). For both methods, ROIs are subsequently eliminated if they do not match the minimum or maximum ROI size, or, if they do not span the minimum number of planes (only in 3D data and when building ROIs in 3D). Implicit maximum distance to dendrite (see section Cleaning of the spine prediction map) is also taken into account. Distances are measured in µm using the user-defined resolution in $x$, $y$, and $z$ of the stack. The settings can be combined if desired.

DeepD3's process of full image segmentation is realized in two applications: A) via a graphical user interface which allows for step-by-step visualization of hyperparameter tuning, and B) via a batch processing function, which can be utilized to segment large amounts of data quickly and in a fully automated fashion once a suited set hyper-parameter settings have been found. Further, DeepD3's classes and functions can be utilized independently in custom code, for example, to modify the 3D ROI building tool or the prediction of dendritic spines and dendrites to the user's needs.

#### Graphical user interface of DeepD3

DeepD3 is accompanied by its built-in graphical user interface (GUI), which allows for fine-tuning of hyper-parameters and visualization of their effects on the image segmentation to the user (Extended Data Figures 8 and 9). The GUI is written in Python 3.7+ and is based on PyQt5 as graphical interface and pyqtgraph for plotting/visualization. We use custom ROI drawing routines to cope with $>$1000 ROIs in a single view. This GUI is provided as a convenience for the user to visualize the stack to be analyzed, the DeepD3 prediction, as well as the 2D and/or 3D ROIs. Moreover, we provide previews of cleaning settings, such that the introduction of artifacts can be reduced. Further, we provide the possibility to get real-time feedback of the 3D ROI building using a dedicated window (Extended Data Figure 9). The main GUI functions comprise (i) the segmentation of dendrites and dendritic spines, (ii) cleaning of the predictions, (iii) 2D ROI building, (iv) 3D ROI building, (v) z-projections of the probability maps. We incorporated the following export options: (i) prediction maps as TIFF files, (ii) ROIs to ImageJ/FIJI format, (iii) ROIs to a folder structure, (iv) ROI map to TIFF file, (v) ROI centroids to CSV file.

A comprehensive video how to install and use DeepD3 can be found on the DeepD3 project website `https://deepd3.forschung.fau.de/`. As a companion to DeepD3, we provide a graphical user interface (GUI) to train a deep neural network according to custom needs (see below).

#### Batch processing function of DeepD3

As an alternative to the inference GUI, DeepD3 comes with the option to batch-process data, once fitting hyper-parameters could be found. This is particularly helpful if large amounts of data with similar image qualities need to be segmented. Using the `batch.py` file allows the user to access a `--help` function that describes the use of the batch script.

#### Protocol for training a custom model

As part of the DeepD3 open framework, we not only allow the inference using pre-trained models from our model zoo, but also allow the users to generate and arrange their own datasets, as well as train their own models. A more detailed description can be found on the DeepD3 Github repository (https://github.com/anki-xyz/DeepD3). Ideally, a dataset is prepared in a dedicated folder. The DeepD3 workflow was tested for universal TIFF images, however, in principle it is not limited to TIFF images. To train your own deep neural network similar as presented in the main body, the following is required:

- Raw or preprocessed microscopy data
- Pixel-wise annotations of the dendritic spines
- Either pixel-wise annotated dendrites or traced dendrites

The two latter points are usually done semi-automatically or manually. In this study, we relied on two convenient ways to generate both: PiPrA (https://github.com/anki-xyz/pipra) for pixel-precise spine annotations and NeuTube for dendrite tracing (https://neutracing.com/download/), see also Figure 1b in the main text for reference. PiPrA already provides a binary segmentation mask as needed for the training algorithm. NeuTube provides SWC files that contain the 3D location of dendritic elements (x, y, z), their given radius (r) and the connection scheme to the previous dendritic element. With this information, we are able to accurately reconstruct the dendrite in 3D. We utilize 3D drawing of spheres that are linear interpolated between defined dendritic elements. The reconstruction also creates a binary segmentation mask similar to PiPrA.

DeepD3 training relies on a custom data loader that conveniently streams tiles from a larger database. To this extent, each dataset needs to be converted to d3data and d3set files (Extended Data Figure 10a). Using our training dataset preparation GUI (Extended Data Figure 10b) one can easily arrange the respective files and select subsets of complex 3D stacks. These subsets can be individually stored as d3data files, which store the part of the original raw data stack, the respective dendritic spine and dendrite segmentation ground-truth, as well as metadata, such as the resolution in x, y and z.

Multiple d3data-files are arranged in a d3set collection (Extended Data Figure 10a,b). This consists of single d3data files and allows for convenient arrangement of training data and easy sharing. The d3set generation is realized using a small GUI element (Extended Data Figure 10b). It allows to select as many d3data files that should be in the training dataset and concatenates those. We recommend creating one d3set each for training and validation data.

Training the DeepD3 architecture itself can be performed by either cloning and using the DeepD3 instance locally or on the Google Colab platform. We provide an example Jupyter notebook for DeepD3 training using the d3sets that have been used to train DeepD3 in the context of this work (see Data Availability).

#### Suggested use of DeepD3

DeepD3 is a versatile tool for quantification of dendrites and dendritic spines. We showed that DeepD3 can be used for various applications, such as spine counting or fluorescence extraction-based analyses. On data similar to the DeepD3 datasets we expect excellent performance. In this work, we showed that even data that is significantly different from the DeepD3 training dataset (species, imaging modality, dye, resolution) DeepD3 performs reliably. However, to optimize the prediction quality we would still recommend to fine-tune DeepD3 pre-trained models to your data or to train the DeepD3 architecture from scratch with your own labeled data (see above instructions). We suggest trying the 32F DeepD3 model that is trained on various resolutions first on your data in the GUI and fine-tune hyperparameters. Depending on its performance, you may elect to try other models from the DeepD3 model zoo or opt to incorporate your own training data. Once a network model and suited hyper-parameters have been found, we recommend using the batch function to quickly and fully automatically segment large quantities of data. N.B. data should thus be processed in batches reflecting similar image qualities (contrast, physical resolution in nm/px), as differences in these image parameters may require different hyperparameter settings for optimal results. In light of the observed amounts of human-to-human variability, we recommend validating performance against manual annotations of multiple human experts on the same data.

### SUPPLEMENTARY DISCUSSION

Imaging of dendritic spines is most frequently performed using two-photon or confocal microscopy. An inherent difficulty of spine imaging using these techniques is resolving fine structures, such as spine necks, small spines or filopodia. As a consequence, spine density is typically underestimated, as some spines are simply missed (Pfeiffer et al., 2018). The limited spatial resolution of these microscopy methods might be one of the underlying factors of the observed levels of inter- and intra-annotator variability. Some regions might simply be too ambiguous to consistently identify spines in. While super-resolution microscopy offers a much clearer assessment of dendrite and spine morphologies, it is challenging to implement in many experimental paradigms. In the future, improvements in image quality during acquisition or in post-processing could contribute considerably to even further improve automated and reliable quantification of dendritic spines.

Some currently available methods of automated spine segmentation have also included options for spine classification. While we provide full segmentation, this type of analysis is in principle easily implemented. Nevertheless, we abstain from this for two reasons. First, the presence of distinct categories of spine morphologies has recently been challenged (Ofer et al., 2021). Second, and more importantly, automated classification seems error-prone at this stage, given the above-mentioned limited spatial resolution most fluorescence microscopes offer (Pfeiffer et al., 2018) and the observed level of variability between manual segmentations (Extended Data Figure 1c).

#### Outlook

With DeepD3, we focus on general applicability across a variety of contexts. Realizing the fast pace of AI, we are sure that DeepD3's technology will be outdated at some point. However, key contributions of the DeepD3 framework is to democratize dendritic spine and dendrite quantification. We offer openly (i) a benchmark dataset annotated by seven raters, (ii) training and evaluation datasets as open data, (iii) the full code to generate own deep neural networks for dendritic spine and dendrite segmentation, (iv) all pre-trained models in an easy ready-to-use fashion, and (v) a GUI and batch processing function for semi- and fully automated spine segmentation. Moreover, we allow the community to contribute to this data repository by (1) uploading neural networks that have been trained using the DeepD3 training procedure and (2) uploading annotated and/or segmented image data using the DeepD3 website (https://deepd3.forschung.fau.de/. We hope that such crowd-sourcing of data and models will continuously improve DeepD3 performance in the future, such that more and more user needs can be met.

In its current iteration, DeepD3 can not match dendritic spines across time points, which is a requirement for the analysis of chronic data. A tool that performs such re-identification of dendritic spines in chronic data is of critical interest to the community. However, due to the sizeable amount of motility dendritic spines show (Bonhoeffer and Yuste, 2002), automating spine re-identification over time-series images is challenging. Mere detection of the spine head is likely not sufficient, as incorrect matches over time could occur. Matching spines using the clustering approach utilized in this manuscript will likely lead to erroneous matches and consequently wrong quantification of spine dynamics. One alternative would be to image the spine neck in sufficient resolution, enabling detection of the base of the spine at the dendrite, which presumably remains unchanged during spine motility. As discussed above, the limited spatial resolution two-photon and conventional confocal imaging offer might not permit precise localization of the spine base and neck. Super-resolution imaging could, in principle, offer such high spatial resolution.

However, certain experimental paradigms are difficult to realize with such imaging methods. While beyond the scope of the current DeepD3 framework, we hope that the challenge of spine re-identification can be overcome in the near future.
